# Fitting bifurcation structure, not voltage traces: Reduced neuron models that preserve physiological parameter dependence

**DOI:** 10.1101/2025.06.14.659682

**Authors:** Louisiane Lemaire, Mahraz Behbood, Jan-Hendrik Schleimer, Susanne Schreiber

**Affiliations:** Institute for Theoretical Biology, Humboldt-Universität zu Berlin, Germany; Bernstein Center for Computational Neuroscience, Humboldt-Universität zu Berlin, Germany; Inria Branch of the University of Montpellier, France

## Abstract

In conductance-based models, spiking-induced ion concentration fluctuations can alter the excitability of single neurons. In particular, in class I models, variations in potassium concentration can induce qualitative changes in the dynamics through a codimension-2 bifurcation known as the saddle-node-loop (SNL). Investigating the implications of such effects at the level of neuronal networks will require computationally efficient single neuron models that still capture ion concentration dynamics realistically. To this end, we propose a method to derive a phenomenological model capturing the coupled extracellular potassium and voltage dynamics of a given class I conductance-based model. Rather than fitting voltage traces, we calibrate a canonical reduced model to the two-parameter bifurcation structure of the target model, with input current and a physiological parameter as coordinates. This preserves the location and type of dynamical transitions as the physiological parameter varies, allowing a single reduced model to capture neuronal dynamics across qualitatively different regimes. The resulting model is an extension of the quadratic integrate-and-fire model, in which extracellular potassium accumulation alters voltage dynamics by increasing the reset voltage. We apply our systematic reduction procedure to the Wang-Buzsáki model. Its phenomenological version exhibits quantitatively comparable dynamics and replicates the reshaping of the phase-response curve associated with the transition from SNIC to HOM spikes at elevated potassium. To illustrate the derived model’s applicability, we perform a preliminary investigation of how changes in potassium concentration influence synchronization in networks.

## Introduction

The biophysical properties of neurons are well-established as critical determinants of both single-cell computation and network behavior. While complex neuron models allow a detailed analysis at the single-cell level, investigation of network dynamics often requires simpler models that can retain essential features of neuronal computation yet ensure computational efficiency [1–3]. This challenge is further intensified when biophysical single-cell properties change dynamically, such as through alterations in the local environment (e.g., temperature fluctuations, changing ionic concentrations) or neuromodulatory effects. To better understand how such modifications influence network dynamics, it is essential to incorporate these physiological changes into simplified neuron models as well. Here, we derive phenomenological neuron models of the integrate-and-fire type that quantitatively capture the dynamics of more complex conductance-based models not only for one specific set of the model’s parameters but also when a physiologically relevant parameter is allowed to vary.

In contrast to the common practice of fitting phenomenological models to spike trains or voltage traces recorded under a single physiological condition [4, 5], we propose to derive phenomenological models by matching a two-dimensional bifurcation structure spanned by the input current as well as an additional physiologically relevant parameter. Here, we demonstrate this approach using the bifurcation diagram of a biologically more complex model (specifically, a conductance-based model including potassium concentration dynamics) as the fitting target. In principle, the same strategy could be applied using a bifurcation diagram reconstructed from experimental data, provided that sufficiently rich measurements are available to characterize the relevant dynamical behavior. Our specific aim is to capture not only the firing rate, but also the bifurcation underlying the onset of regular firing at threshold, including switches in bifurcation type as the physiological parameter varies. While we focus here on the influence of extracellular potassium, the approach is equally applicable to other biological parameters.

In the processing of information within single cells, the dynamics of action-potential generation play an important role. Although influenced by numerous cellular characteristics, qualitative differences between regular action-potential firing (with all-or-none pulses at threshold) can arise from three distinct mechanisms. Their classification coincides but is not entirely identical with the excitability classes I and II described by Hodgkin (1948) [6] and observed earlier by Arvanitaki (1938) [7], which are based on the frequency of firing at spike onset (with arbitrarily low frequencies for class I and an immediate jump to higher frequencies for class II). These two classes are associated with distinct mathematical bifurcations that underlie the onset of regular spiking: the saddle-node-on-invariant circle (SNIC) bifurcation is the canonical class-I onset mechanism, while class II is typically associated with the subcritical Hopf bifurcation. Another bifurcation, the saddle-homoclinic orbit (HOM) bifurcation, shares the arbitrarily low onset frequencies of class I while differing in other properties (see Izhikevich [8] for details). The type of bifurcation is a critical determinant of the network state. For instance, neurons with SNIC dynamics tend to produce asynchronous activity when weakly coupled within inhibitory networks. In contrast, the transition to HOM dynamics in the same network configuration results in synchronous activity [9].

The transition from SNIC to HOM is of particular interest, as it is ubiquitous in the underlying mathematical framework (despite a quantitative dependence of the switching point on other parameters) and has the potential to induce dramatic changes in network behavior [10, 11]. One well-established parameter that causes this transition is the concentration of extracellular potassium. It is directly affected by neuronal spiking. While the outflow of K^+^ ions at each spike is typically compensated for by mechanisms such as Na^+^/K^+^-ATPases and glial buffering, these may not suffice to prevent transient changes during intense activity [12]. Such elevations alter the neuron’s excitability, forming a feedback loop that can create rich dynamics [13]. Moreover, abnormally elevated levels of extracellular potassium are involved in paroxysmal pathological activities that are specific to epilepsy (seizures) and migraine with aura (spreading depolarization) [14].

To include the effect of extracellular potassium concentration in simpler, network-compatible phenomenological models, we start from a conductance-based model including the potassium concentration dynamics and fit a simpler, integrate-and-fire-type model equipped with a potassium variable to this conductance-based model, aiming to capture both firing rates and the two-dimensional bifurcation structure with input current and extracellular potassium concentration as parametric dimensions. Fitting the bifurcation structure of the reduced model near the SNL bifurcation accurately captures the conductance-based model behavior under a range of ionic concentrations, associated with qualitative changes in excitability. This permits the study of transitions between those different regimes as potassium evolves dynamically. Accordingly, the phenomenological model captures a transition from SNIC to HOM firing with increases in extracellular potassium.

Overall, our contribution is twofold: a minimal model of potassium-dependent excitability and, more generally, a bifurcation-guided procedure for embedding physiologically meaningful parameter dependence into canonical reduced neuron models. In the following, we first demonstrate how to quantitatively derive the integrate-and-fire type version of a given class I conductance-based model. We then compare, at the single-neuron level, the dynamics of the two models and their phase-response curves. Finally, we investigate the effect of potassium on neuronal synchronization at the network level.

## Results

### Reducing a class I conductance-based model to a QIF neuron with dynamic [K^+^]_o_

We demonstrate our procedure on a Wang-Buzsáki model embedded in an extracellular space with dynamic potassium concentration (Fig. 1a) [13, 15], hereafter referred to as the target model. We reduce it to a model of type quadratic integrate-and-fire (QIF), which allows us to capture the normal forms of both SNIC and HOM bifurcations, as well as the transition between them at the so-called saddle-node-loop (SNL) point via the reset voltage (see Fig. S1). Reset values below the voltage at the saddle-node bifurcation of the QIF correspond to SNIC onsets, while larger values correspond to HOM onsets. We carefully choose parameter values of the QIF system and incorporate potassium dependencies to match the homoclinic, saddle-node, and SNIC bifurcation branches of the target model near the SNL transition. In particular, potassium concentration feeds back on the reset voltage, inducing the SNL. An additional reset rule, which increments potassium concentration by a small amount at each spike, mimics the flow of ions through voltage-gated potassium channels (Fig. 1b). Including an electrogenic Na^+^/K^+^-ATPase complements the potassium dynamics. The resulting reduced model is, like the original, a slow-fast system where slow changes in potassium drive the neuron across different excitability regimes.

**FIG. 1:**
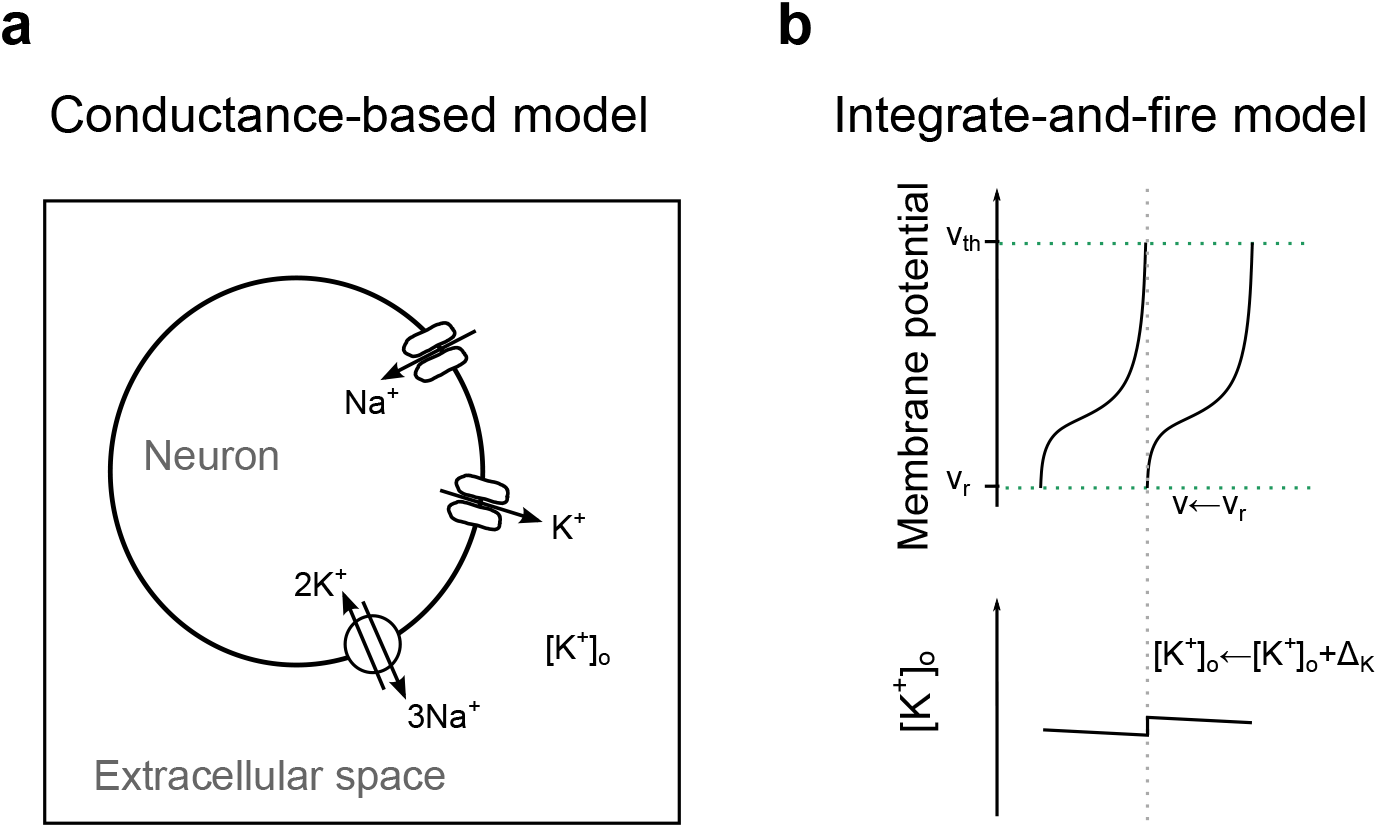
Schematic representation of the two models: the target model (a) and its corresponding phenomenological version (b). **(a)** Conductance-based model of excitability class I with dynamic extracellular potassium concentration, equipped with the usual Na^+^ and K^+^ voltage-gated channels and with an electrogenic Na^+^/K^+^ ATPase (diagram reproduced from [13]). **(b)** Quadratic integrate-and-fire (QIF) model with dynamic extracellular potassium concentration, equipped with an electrogenic Na^+^/K^+^ ATPase. Each spike (i.e., when the voltage *v* exceeds a threshold *v*_th_) triggers two reset rules: as in the classical QIF model the voltage is reset to a value *v*_r_, and the extracellular potassium concentration is additionally incremented by a small amount Δ_K_.

#### Original model

For convenience, we reproduce in “Methods” the definition of the extended Wang-Buzsáki model [13] (System (4)), which we use as an example to illustrate the reduction procedure.

Relying on the difference in timescale between [K^+^]_o_ and the other variables, the fast subsystem of System (4) can be defined as Eqs. (4a) to (4c), where [K^+^]_o_ is a parameter. At physiological potassium concentrations, spike onset is mediated by a SNIC bifurcation (e.g., when [K^+^]_o_ = 5 mM, see Fig. 2a), while at larger concentrations it is mediated by a homoclinic bifurcation (e.g., when [K^+^]_o_ = 12 mM, see Fig. 2b) [13]. Fig. 3a shows the SNL bifurcation organizing this transition.

**FIG. 2:**
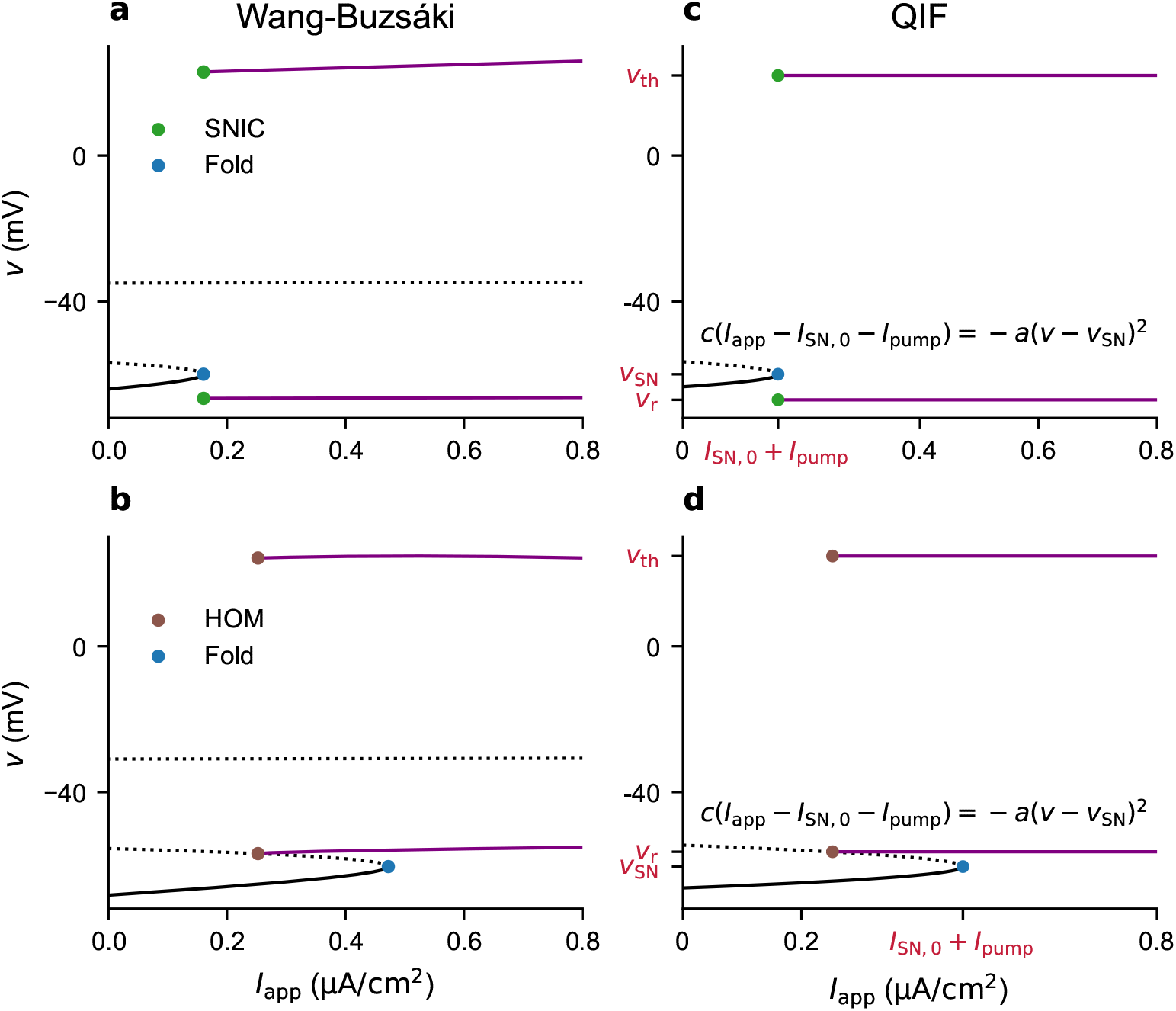
Matching the bifurcation structure of the target model with the QIF model, at fixed values of [K^+^]_o_. **(a-b)** Bifurcation diagrams of the fast subsystem of the Wang-Buzsáki model (Eqs. (4a) to (4c)) as a function of the applied current, featuring SNIC spike onset when [K^+^]_o_ = 5 mM (a) and homoclinic onset when [K^+^]_o_ = 12 mM (b). **(c-d)** Bifurcation diagrams of the model defined in Eqs. (1a) and (1b), with parameter values chosen to mimic the diagrams in (a) or (b), respectively. Limit cycles are shown in purple and fixed points in black, with solid lines for stable branches and dotted lines for unstable branches.

**FIG. 3:**
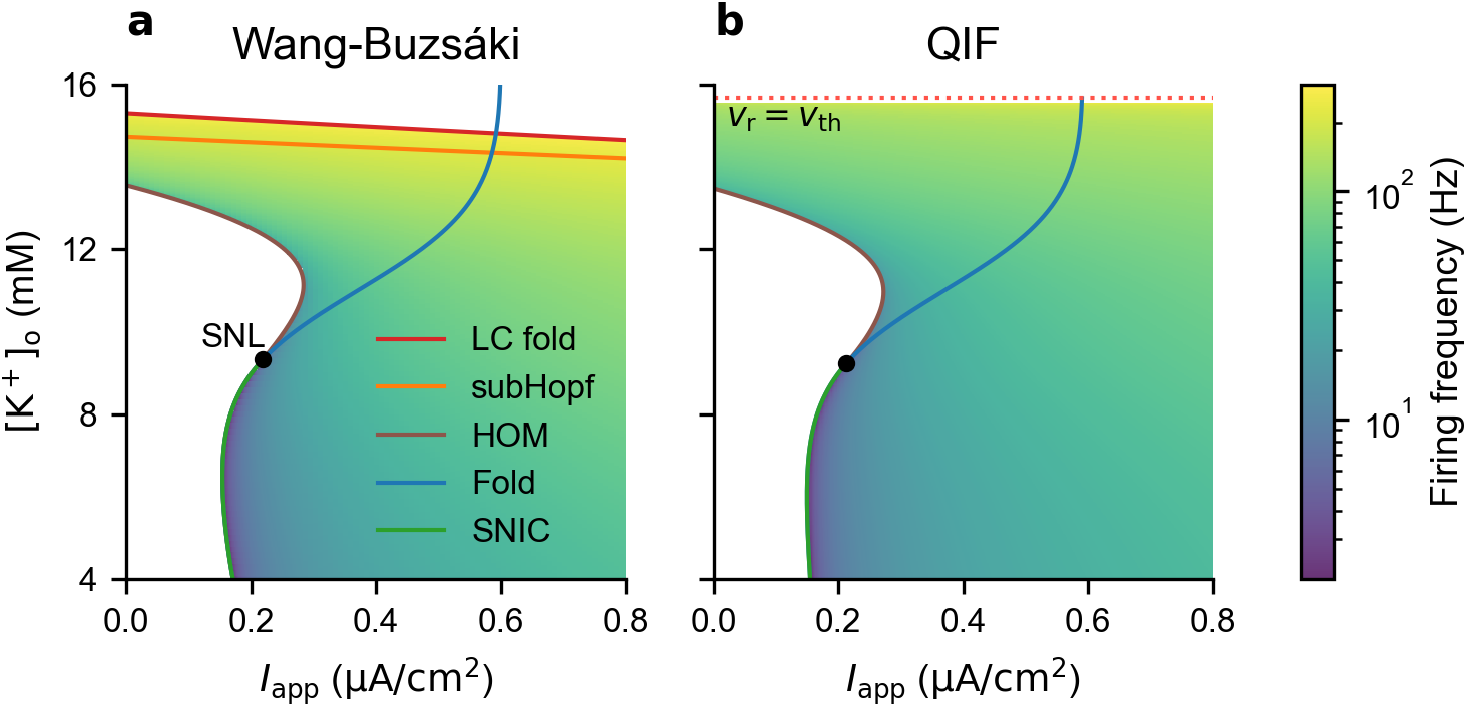
Bifurcation structure and frequency landscape near the SNL bifurcation. **(a)** Two-parameter bifurcation diagram [K^+^]_o_ versus applied current *I*_app_ of the fast subsystem of the Wang-Buzsáki model (Eqs. (4a) to (4c)) and frequency of the limit cycles. **(b)** Same as (a), for the QIF model (Eqs. (1a) and (1b)). Note that in this model, the firing frequency goes to infinity as *v*_r_ approaches *v*_th_. Here, we only show the frequencies up to a certain distance between those parameters.

When the full system slowly follows a branch of stable *fixed points* of the fast subsystem’s bifurcation diagram, i.e., when the neuron is quiescent, the dynamics of [K^+^]_o_ can be studied using the standard slow subsystem (System 6). When the full system slowly follows a branch of stable *limit cycles*, i.e., when the neuron spikes, each spike causes a rapid increment of [K^+^]_o_ (note the presence of *v* in Eq. (4d)). However, these increments are of small amplitude, and the dynamics of [K^+^]_o_ remains slow on average (see already Fig. 6c). As is customary in such cases [16–18], to isolate the slow component of the dynamics, the averaging method can be applied to define the averaged slow subsystem (System 7). The strategy consists in averaging the derivative of the slow variable [K^+^]_o_ over one period (i.e., one spike cycle), for each member of the family of fast subsystem stable limit cycles.

#### Step 1: QIF version of the Wang-Buzsáki model for a constant [K^+^]_o_

We consider the following QIF model:

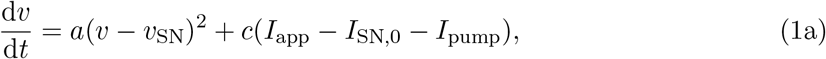

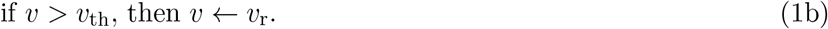

Its branch of fixed points, defined by *c*(*I*_app_ − *I*_SN,0_ − *I*_pump_) = −*a*(*v* − *v*_SN_)^2^, folds at (*I*_app_ = *I*_SN,0_ + *I*_pump_, *v* = *v*_SN_). Stable limit cycles emerge via a SNIC bifurcation when the reset voltage *v*_r_ is smaller than *v*_SN_ (Fig. 2c), while they emerge via a homoclinic bifurcation at 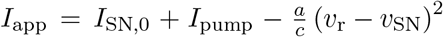 when *v*_r_ *> v*_SN_ (Fig. 2d), with a SNL bifurcation at *v*_r_ = *v*_SN_. Let us first determine, for a given [K^+^]_o_, parameter values of System (1) such that its bifurcation diagram with respect to the applied current closely resembles that of the fast subsystem of the target model.

Note that in the target model (System (4)), *I*_pump_ depends only on potassium (Eq. (5d)). We can therefore treat this current as a potassium-dependent negative contribution to the applied current. In other words, when [K^+^]_o_ is fixed, *I*_pump_ only horizontally translates bifurcation diagrams with respect to *I*_app_, such as those shown in Fig. 2. The parameter derivation below is performed in the absence of the pump current. *I*_pump_ can be added a posteriori, taken to be the same as in the target model (Eq. (5d)).

*I*_SN,0_ is the value of applied current *I*_app_ at which the saddle-node bifurcation of interest occurs in the fast subsystem of the target model, in the absence of pump current. *v*_SN_ is the membrane potential at the saddle-node. We take *a* and *c* to be the quadratic normal form coefficients for the saddle-node bifurcation. They are defined as (see, for example, [19]):

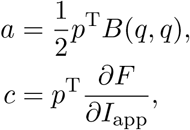

where *p* and *q* are the normalized left and right eigenvectors of the Jacobian matrix and *B* the Hessian quadratic form of the vector field 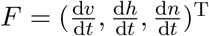 at the saddle-node:

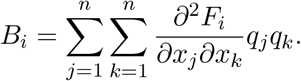

Since the bias term *c* is close to 1 cm^2^ µF^−1^, we set it to this value for simplicity.

Then, if [K^+^]_o_ is larger than the value at the SNL bifurcation in the fast subsystem of the target model [K^+^]_o,SNL_, we choose *v*_r_ such that the homoclinic bifurcation in the QIF model takes place at the same value of applied current:

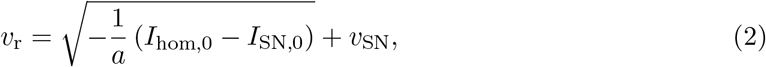

where *I*_hom,0_ is the value of *I*_app_ at which the homoclinic bifurcation occurs in the fast subsystem of the target model, in the absence of pump current. Note that *I*_hom_ = *I*_hom,0_ + *I*_pump_ is the value of *I*_app_ at which the homoclinic bifurcation occurs in the presence of a pump current.

When [K^+^]_o_ *<* [K^+^]_o,SNL_ (i.e., in the SNIC case), we have more flexibility in the choice of the reset voltage. *v*_r_ should be smaller than *v*_SN_ to obtain a SNIC bifurcation. *v*_r_ should also be reasonably close to the limit cycles’ minimum voltage in the target model. We choose to define *v*_r_ in the SNIC case by extrapolating the linear fit of the values of *v*_r_ assigned using Eq. (2) in the homoclinic case (i.e., when [K^+^]_o_ *>* [K^+^]_o,SNL_) to lower values of [K^+^]_o_ (see Fig. 4d). This satisfies the two criteria above.

**FIG. 4:**
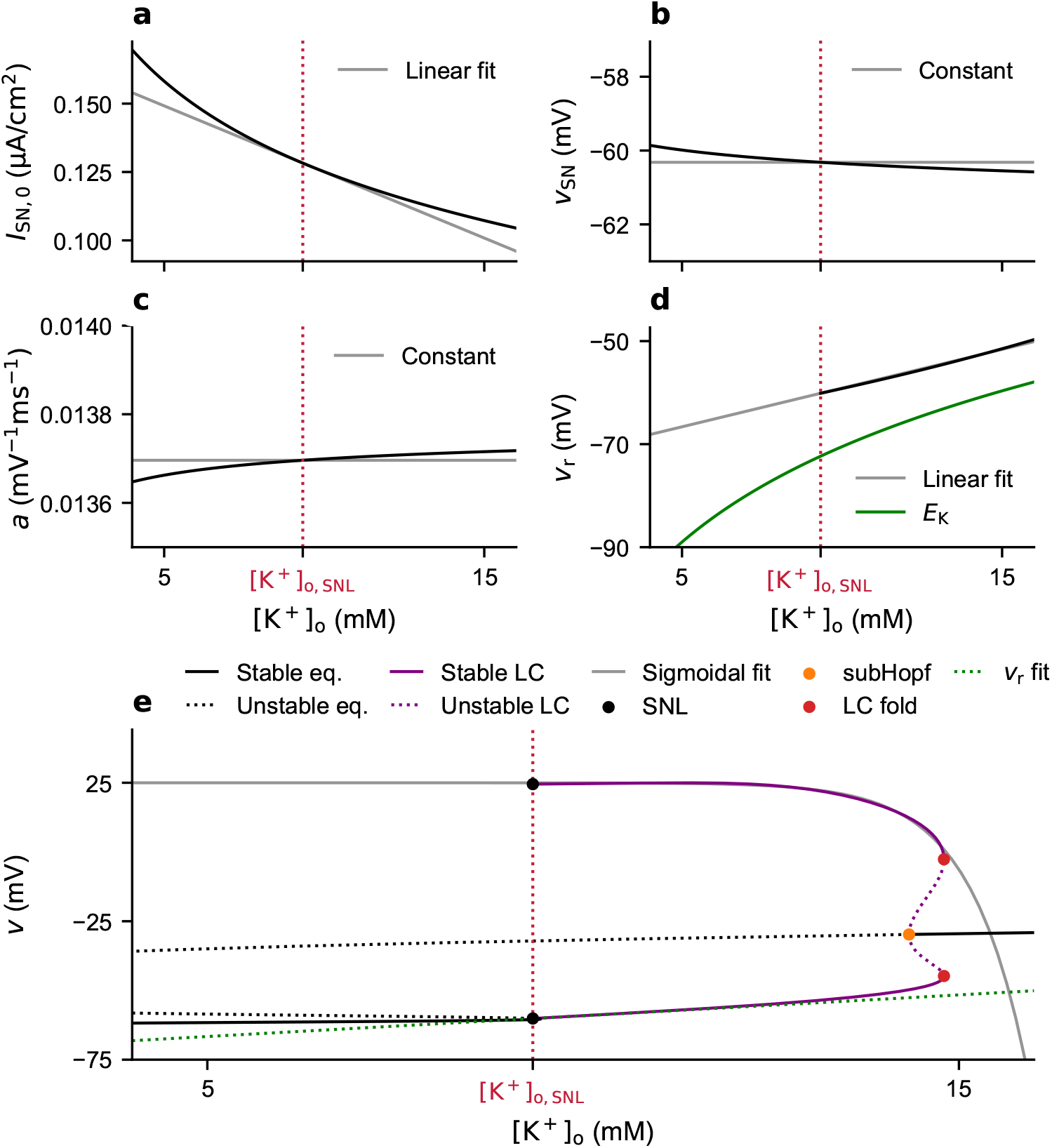
Parameters of the QIF model (Eqs. (1a) and (1b)) depending on the potassium concentration. **(a-d)** Derivation of the parameters *I*_SN,0_, *v*_SN_, *a* and *v*_r_. Note that the increase of the reset voltage *v*_r_ with [K^+^]_o_ reflects the increase of the potassium reversal potential *E*_K_ in the conductance-based model (panel (d)). **(e)** Bifurcation diagram of the fast subsystem of the target model (Eqs. (4a) to (4c)) with respect to [K^+^]_o_, when *I*_app_ = *I*_app,SNL_ (i.e., the value of applied current at which the SNL bifurcation occurs) and *I*_pump_ = 0 µA cm^−2^. In this panel, we have reproduced the linear fit of the reset voltage *v*_*r*_ shown in panel (d), for visual comparison with the minimal voltage values of the limit cycles.

Finally, the threshold voltage *v*_th_ will allow us to approximate the limit cycles’ maximum voltage.

#### Step 2: QIF version of the Wang-Buzsáki model with dynamic [K^+^]_o_

We now compute parameter values of System (1) for a range of [K^+^]_o_, and depending on how strongly they vary with potassium, either set them to a constant or replace them with a function of [K^+^]_o_.

Fig. 4a-d shows *I*_SN,0_, *v*_SN_ and *a* for [K^+^]_o_ between 4 and 16 mM and *v*_r_ only above [K^+^]_o,SNL_. We notice that *v*_SN_ and the normal form coefficient *a* are relatively conserved in this interval. We fix them to their value at [K^+^]_o,SNL_. For *I*_SN,0_ and *v*_r_ we use linear fits, constrained at [K^+^]_o,SNL_, that we extrapolate to lower concentrations in the case of *v*_r_. In conductance-based models, higher [K^+^]_o_ weakens the potassium concentration gradient, thereby raising *E*_K_. The sodium and potassium reversal potentials typically bound voltage oscillations during spiking (see, for example, [20]), so the minimum voltage of spikes is expected to be higher when [K^+^]_o_ is larger. This effect is reflected in the QIF model via the increase of the reset voltage *v*_r_ (Fig. 4d).

For the threshold *v*_th_, we use a sigmoid that we extrapolate to lower concentrations, to fit the peak voltage of the limit cycles in the bifurcation diagram with respect to [K^+^]_o_, when *I*_app_ = *I*_app,SNL_ and in the absence of pump (Fig. 4e). This enables us to capture the shrinkage in spike amplitude that precedes the fold of limit cycles bifurcation. Note that, following the SNL bifurcation, the peak voltage increases slightly with [K^+^]_o_, by less than 1 mV. We do not include this increase in the QIF model.

We incorporate these functions of [K^+^]_o_ in System (1), in which [K^+^]_o_ now appears as a parameter. Its bifurcation structure with respect to [K^+^]_o_ and the applied current *I*_app_ closely resembles that of the fast subsystem of the original model (see already Fig. 3). A summary of the reduction procedure up to this stage is provided in the form of an algorithm in Table I. An illustration of this procedure using temperature instead of potassium concentration is shown in Fig. S2. Returning to the potassium example, we then include potassium dynamics to obtain the following model:

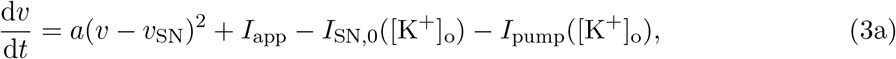

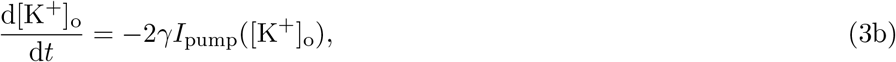

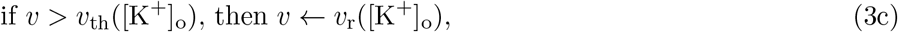

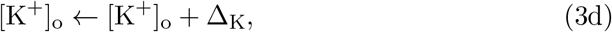

with

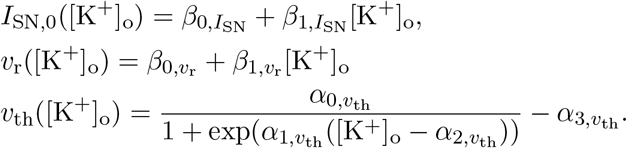

**TABLE I:**
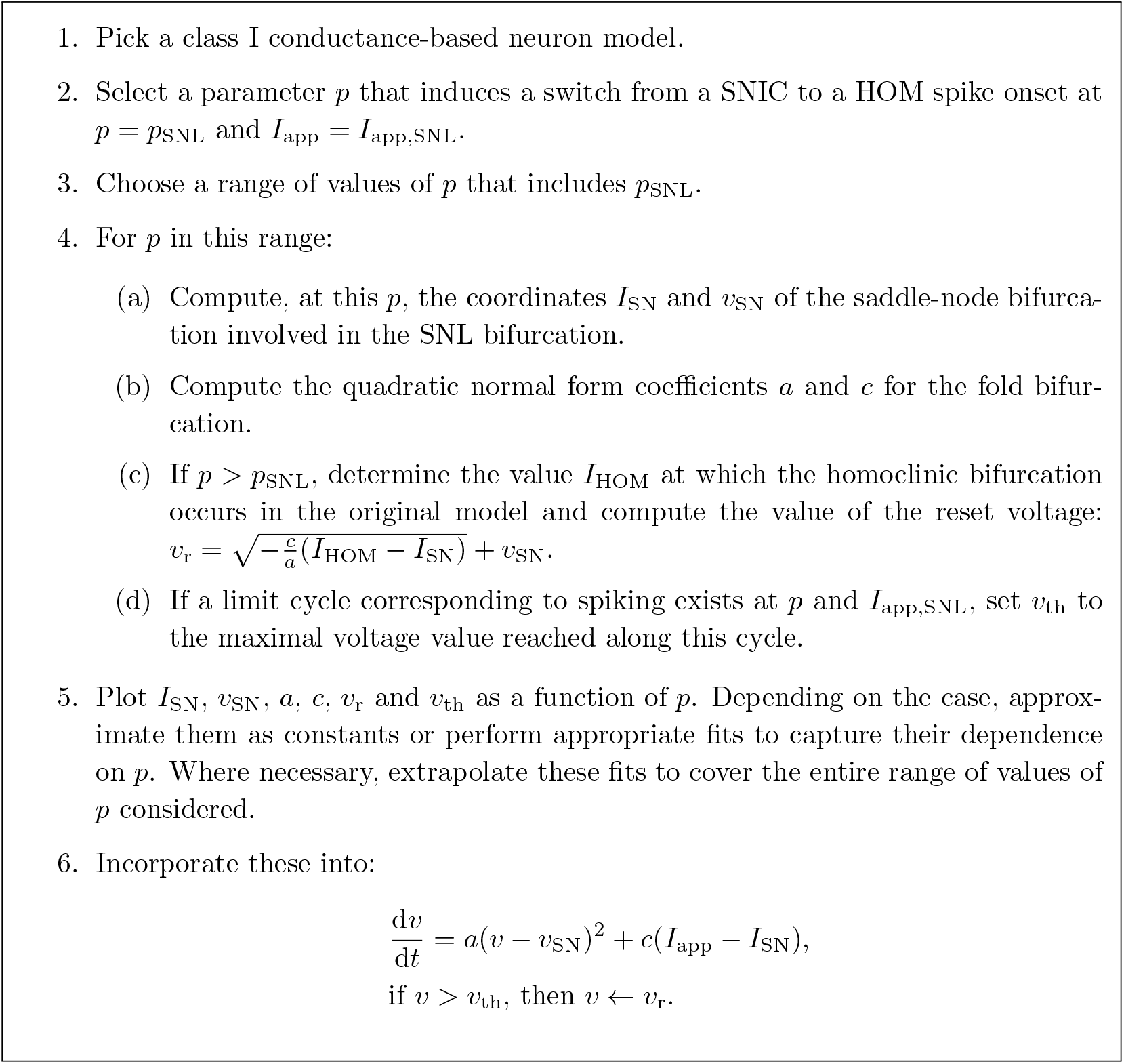
Steps to derive a quadratic integrate-and-fire model with dependence on a biophysical parameter, by fitting the bifurcation structure of a class I conductance-based neuron model near an SNL bifurcation induced by this parameter. Note that, in this paper, we performed these reduction steps in the absence of pump current in the original model, and added this current a posteriori to the reduced model.

The evolution of the extracellular potassium concentration depends, on the one hand, on uptake by the Na^+^/K^+^-ATPase (Eq. (3b), where *I*_pump_ is given in Eq. (5d)), and on the other hand, on increments at each action potential (Eq. (3d)). Note that the reset rule Eq. (3d) captures the dependency of potassium dynamics on *v* in the original model (Eq. (4d)). When the target model is Wang-Buzsáki, the parameter values of the corresponding QIF version in the form of (3) are listed in Table II.

**TABLE II:**
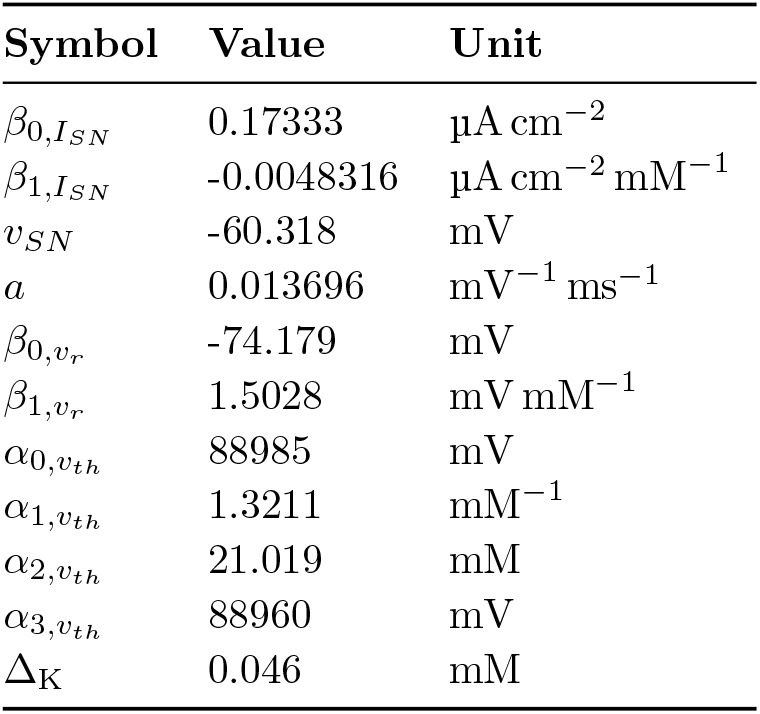
Parameter values of the QIF model (3) to match the bifurcation structure of the fast subsystem of the Wang-Buzsáki model (4) with respect to [K^+^]_o_. The values of the parameters *γ, I*_pump,max_, *k* and K_eq_ are unchanged compared to Table IV.

The fast subsystem of this model consists of Eqs. (3a) and (3c), with [K^+^]_o_ treated as a parameter. Its bifurcation diagram with respect to input and potassium, and the frequency of the limit cycles, can easily be calculated analytically (see “Methods”). We can see in Fig. 3 that, with our reduction procedure, we achieve a very close match of the bifurcation branches emanating from the SNL bifurcation in the fast subsystem of the target model. In Wang-Buzsáki, the transition to depolarization block at larger potassium concentrations is mediated by a fold of limit cycles and a subcritical Hopf bifurcation (panel a). The dimensionality of the QIF fast subsystem does not allow for such bifurcations. Instead, we artificially obtain a depolarization block by silencing the dynamics when the threshold voltage *v*_th_ drops to the reset level (see gray and green curves in Fig. 4e). Note that, although the firing frequency in the fast subsystem grows to infinity as [K^+^]_o_ approaches this point, this does not effectively occur in the full system since [K^+^]_o_ is incremented by a constant amount with each spike.

The standard slow subsystem of System (3), used to study the dynamics of [K^+^]_o_ along stable branches of fixed points of the fast subsystem, is given by

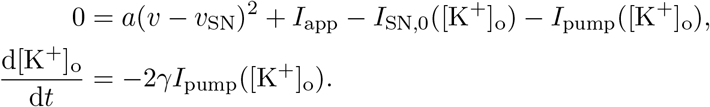

As in the original model, along stable branches of limit cycles of the fast subsystem, the dynamics of [K^+^]_o_ includes a fast component, causing rapid increments at each spike (here, instantaneous jumps via the reset rule Eq. (3d)). The jump size Δ_K_ is chosen to match the potassium increment per spike in the original model and is therefore small (see already Fig. 6h). Thus, the averaging method can still be employed (see, for example, [8]). The averaged slow subsystem of System (3) is then given by

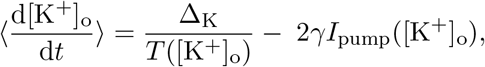

where *T* is the period of the limit cycles.

### Our phenomenological neuron model closely captures the dynamics of the original conductance-based version

Under identical stimulation, the behavior of the phenomenological model resulting from this bifurcation diagram fitting procedure is very similar to that of the target model. In the same way as the original version, it can produce regular tonic firing (Fig. S4), as well as richer firing patterns. We illustrate this point with three scenarios typically occurring in class I conductance-based models [13, 16, 21], which can notably be obtained in the Wang-Buzsáki model (see overview in Fig. 5). They share the common feature of involving switches in firing regimes, caused by an activity-mediated accumulation of extracellular potassium. Those switches rely on: (i) the different regimes associated with each region of their common bifurcation diagram and (ii) slow (on average) changes in [K^+^]_o_. Although the time traces are not exactly the same, we will see that the two models are quantitatively comparable.

**FIG. 5:**
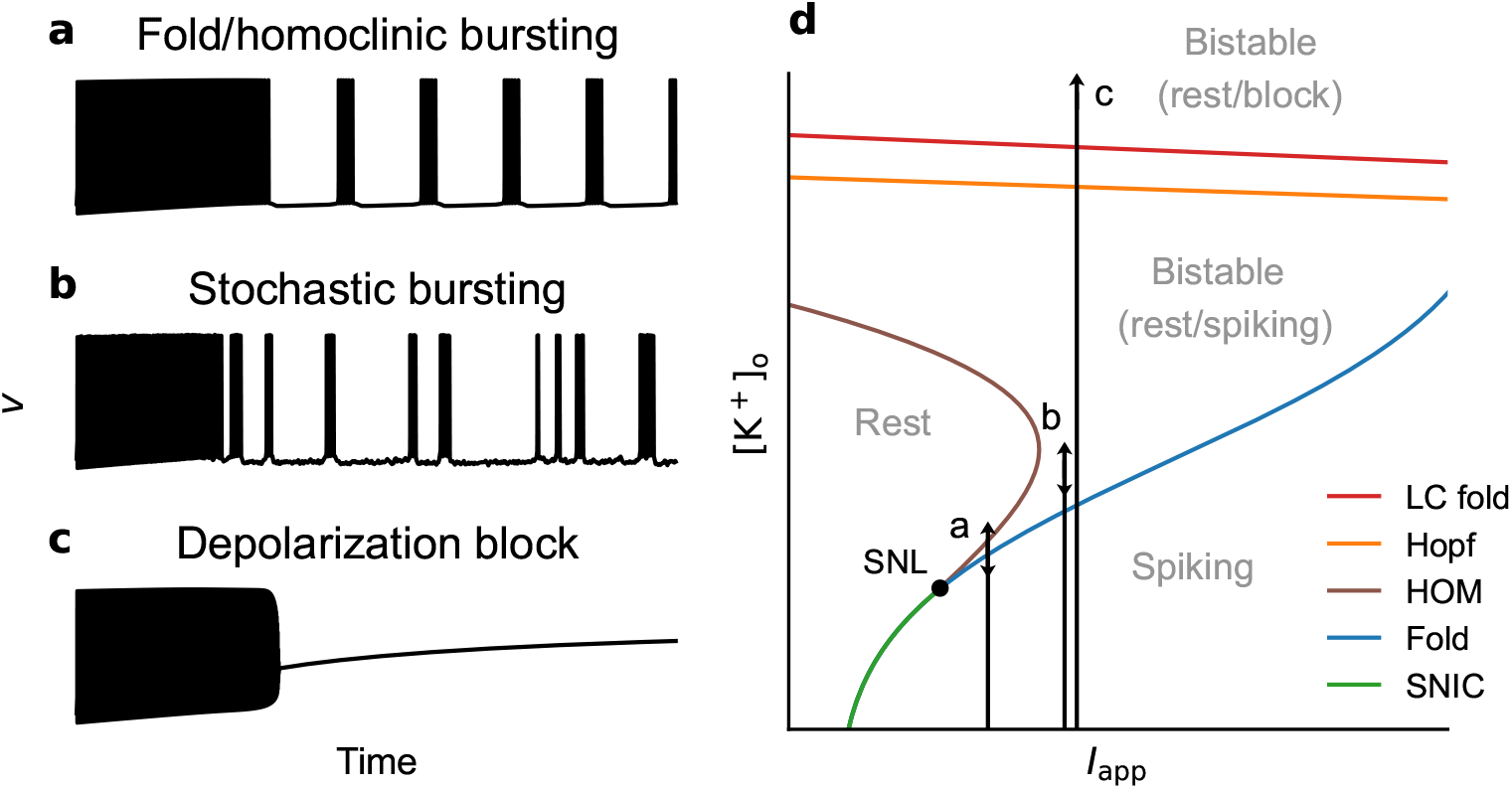
Three scenarios of activity-mediated switches in firing regimes that were reported in class I conductance-based models. **(a-c)** Voltage traces, in the case of the Wang-Buzsáki model. **(d)** Schematic representation of trajectories in the [K^+^]_o_ versus applied current bifurcation diagram through regions associated with various activity regimes.

#### Scenario 1: fold/homoclinic bursting

We first consider the hysteresis loop bursting mechanism described in [13], which relies on the interplay between slow changes in extracellular potassium concentration and voltage dynamics. It requires only fast-inactivating sodium channels, delayed rectifier potassium channels, and an indispensable electrogenic pump, without the need for any slow ion channels.

Starting from a physiological potassium concentration ([K^+^]_o_ = 5 mM [22]), we stimulate the Wang-Buzsáki neuron with a constant deterministic applied current (*I*_app_ = 0.25 µA cm^−2^). This corresponds to an initial condition in the spiking region (Fig. 6a), i.e., where the only attractor is a stable limit cycle with large voltage amplitude (Fig. S3a). There, potassium release through voltage-gated channels outweighs the effect of the pump on average over the spike cycles, leading to a progressive accumulation of extracellular potassium (first part of Fig. 6b).

**FIG. 6:**
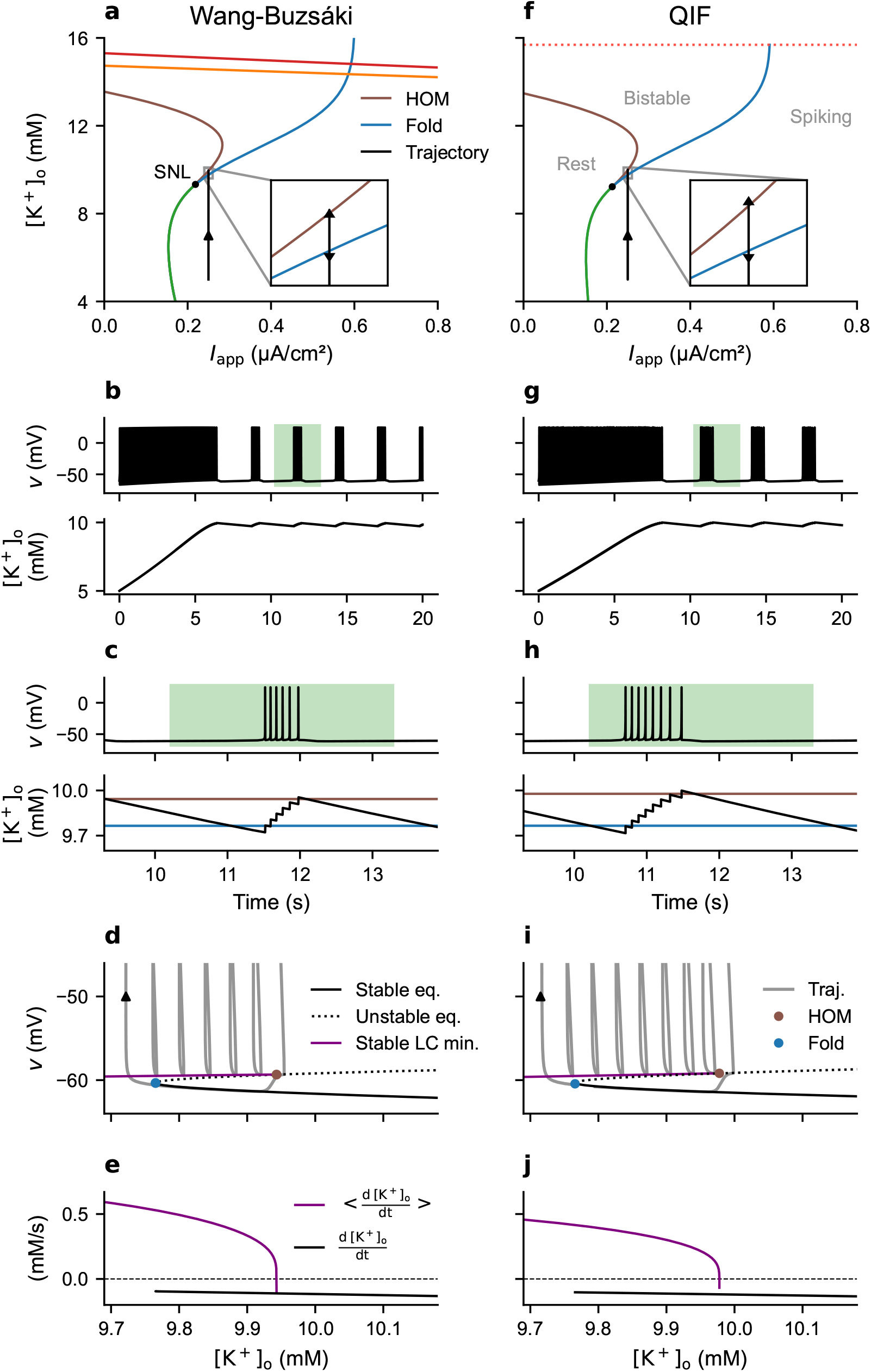
Fold/homoclinic bursting scenario (deterministic input: *I*_app_ = 0.25 µA cm^−2^). For the Wang-Buzsáki model: **(a)** Bursting trajectory shown on the 2-parameter bifurcation diagram [K^+^]_o_ versus *I*_app_ of the fast subsystem. **(b)** Voltage and [K^+^]_o_ time traces. **(c)** Enlargement of the burst highlighted in panel (b). **(d)** Bursting trajectory superimposed onto the 1-parameter bifurcation diagram of the fast subsystem with respect to [K^+^]_o_, zooming in on lower voltage values for better visibility. **(e)** Time derivative of the averaged and standard slow subsystems. For the quadratic integrate-and-fire model: **(f-j)** Same as (a-e).

When the system arrives in the area enlarged in Fig. 6a, where the bistable region is bounded from below (lower [K^+^]_o_) by the fold bifurcation and from above (higher [K^+^]_o_) by the homoclinic bifurcation, it begins to burst (second part of Fig. 6b). For completeness, note that this configuration is only a necessary condition for the bursting behavior to occur (for more details, see [13]). During each spiking phase, the system follows the branch of stable limit cycles (Fig. 6d), in the course of which potassium increases overall (in the averaged slow subsystem 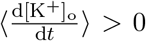, see purple curve in Fig. 6e) because the increments at each action potential are not fully compensated by the pump (Fig. 6c). When it reaches the homoclinic bifurcation, the system switches to the branch of stable equilibria, initiating a quiescent phase. In the absence of spiking, the pump can restore potassium concentration down to the fold bifurcation (in the standard slow subsystem 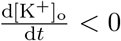, see the black curve in Fig. 6e), at which point the system jumps back to the stable limit cycle branch. This marks the start of a new bursting cycle.

Starting from the same initial voltage and potassium concentration, and stimulated with the same input, the integrate-and-fire version of the model successfully captures this bursting behavior (Fig. 6f-j). The underlying mechanism is the same as in the original model, with fold and homoclinic dynamical bifurcations governing the onset and termination of bursts. The small discrepancy between the two models in the number of spikes per burst and interspike interval is due to the divergence in the frequency of limit cycles (Fig. S3d) and slightly different location of the bifurcations in relation to potassium levels.

We should point out here that, as explained in Section 9.2.3 of [8], the reduction of the dynamics to the averaged slow subsystem is not valid in the close vicinity of the homoclinic bifurcation. Therefore, the intersection between the x-axis and the time derivative of the averaged slow subsystem (purple curve), visible in Fig. 6e and j, does not imply the existence of a stable limit cycle corresponding to tonic firing in the full system.

For this first scenario, in accordance with [13], we adopted a configuration where the potassium concentration rises until the neuron starts bursting. However, note that simulations can also be generated in which the neuron remains in the spiking region, for example, by implementing a buffering of the extracellular potassium. This can conveniently be done in both original and reduced versions of the model by introducing a term in the potassium equation. As a result, when the buffering is sufficiently strong, potassium stabilizes within the spiking region at a concentration corresponding to a fixed point of the averaged slow subsystem, which in this case equates to a limit cycle of the full system (Fig. S4).

#### Scenario 2: stochastic bursting

We now examine a switch in firing regime identified in a version of the Traub-Miles model [23] (a class I conductance-based model) accounting for potassium concentration dynamics. By introducing a noise component in the input, Contreras et al. obtained simulations showing that, after a period of tonic firing, the neuron switches to stochastic bursting, a regime where firing is intermittently interrupted [21]. It is possible to reproduce this behavior with the Wang-Buzsáki model.

We stimulate the neuron with colored noise (see Eq. (9) in “Methods”) of mean input *I*_app_ = 0.3 µA cm^−2^, chosen sufficiently large not to interfere with the fold/homoclinic bursting behavior (Fig. 7a). Initially, like in the previous scenario, the neuron spikes periodically, causing an accumulation of extracellular potassium (first part of Fig. 7b). When the system reaches the bistable region, i.e., when [K^+^]_o_ crosses the fold bifurcation of the fast subsystem, the firing pattern switches to stochastic bursting (second part of Fig. 7b). There, the noisy input triggers jumps between the two attractors: the stable fixed point corresponding to a resting state and the stable limit cycle corresponding to tonic firing (Fig. 7d).

**FIG. 7:**
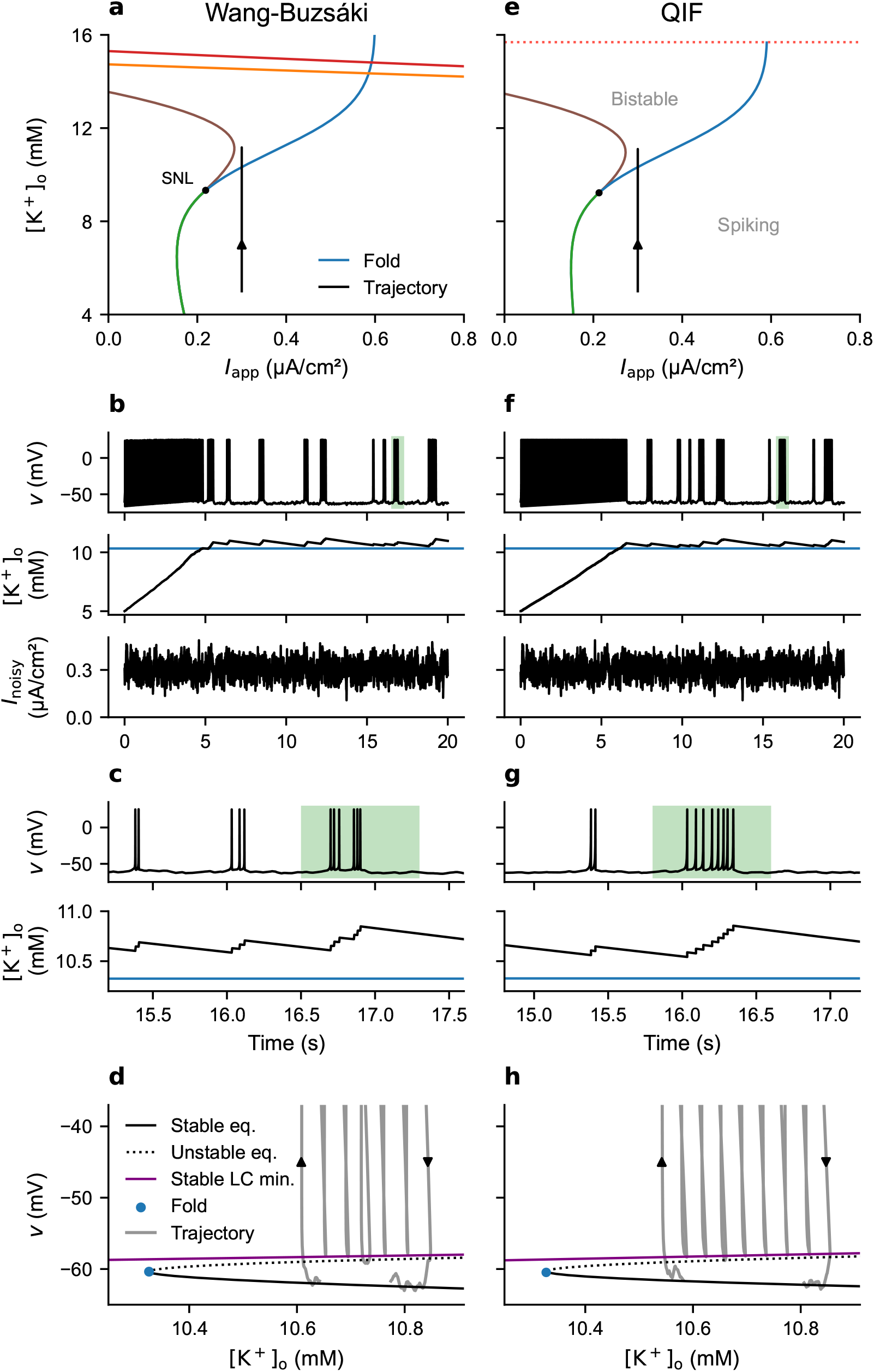
Stochastic bursting scenario (noisy input, with mean *I*_app_ = 0.3 µA cm^−2^). For the Wang-Buzsáki model: **(a)** Trajectory shown on the 2-parameter bifurcation diagram [K^+^]_o_ versus *I*_app_ of the fast subsystem. **(b)** Voltage, [K^+^]_o_ and applied current time traces. **(c)** Enlargement of the burst highlighted in panel (b). **(d)** Part of the trajectory corresponding to the burst highlighted in panels (b) and (f), superimposed onto the 1-parameter bifurcation diagram of the fast subsystem with respect to [K^+^]_o_. For the quadratic integrate-and-fire model: **(e-h)** Same as (a-d).

As expected from their common bifurcation structure, when confronted with the same stimulus (same realization of the stochastic process) as the target model, the integrate-and-fire version behaves very similarly (Fig. 7e-h).

Compared to the deterministic fold/homoclinic bursting, for which burst initiation and termination occur only at the boundaries of the bistable region, in this scenario the noise can induce switches between attractors within the bistable region (Fig. 7c-d,g-h).

On a side note, we observe an interruption of about 80 ms in the middle of the burst highlighted in Fig. 7c, during which the voltage is more depolarized than during interburst intervals. Instead, the system remains near the repelling slow manifold (the saddle branch in Fig. 7d). The transition from following an attracting manifold during the burst to evolving near a repulsive one is a canard-like effect, here induced by noise. Similar canard trajectories can be obtained in a deterministic way, see for example [24].

#### Scenario 3: transition to depolarization block

The last scenario we consider is the transition to depolarization block, which results in transient spike inactivation. This phenomenon is a pathological hallmark: it is the signature of cortical spreading depolarization at the single-neuron level and might also be involved in epileptiform activity [25]. The induction of a depolarization block, in the case where it is caused by an accumulation of extracellular potassium resulting from neuronal firing, has been extensively studied in conductance-based models with dynamic ion concentrations (e.g., [16, 21, 26–29]). We simulate it here with the Wang-Buzsáki model.

To achieve this, we use the same setup as for the stochastic bursting scenario, but without the noise component in the stimulus. As a result, in the bistable region, the system remains on the branch of stable limit cycles, allowing the extracellular potassium concentration to build up without interruption. When [K^+^]_o_ exceeds the concentration corresponding to a fold of limit cycles of the fast subsystem, the neuron enters depolarization block (Fig. 8a-d). In comparison, in the stochastic bursting scenario, frequent pauses in firing due to the presence of noise make it easier for the regulatory mechanisms—in our model, the Na^+^/K^+^-ATPase—to maintain [K^+^]_o_ in the bistable region, preventing the pathological block.

**FIG. 8:**
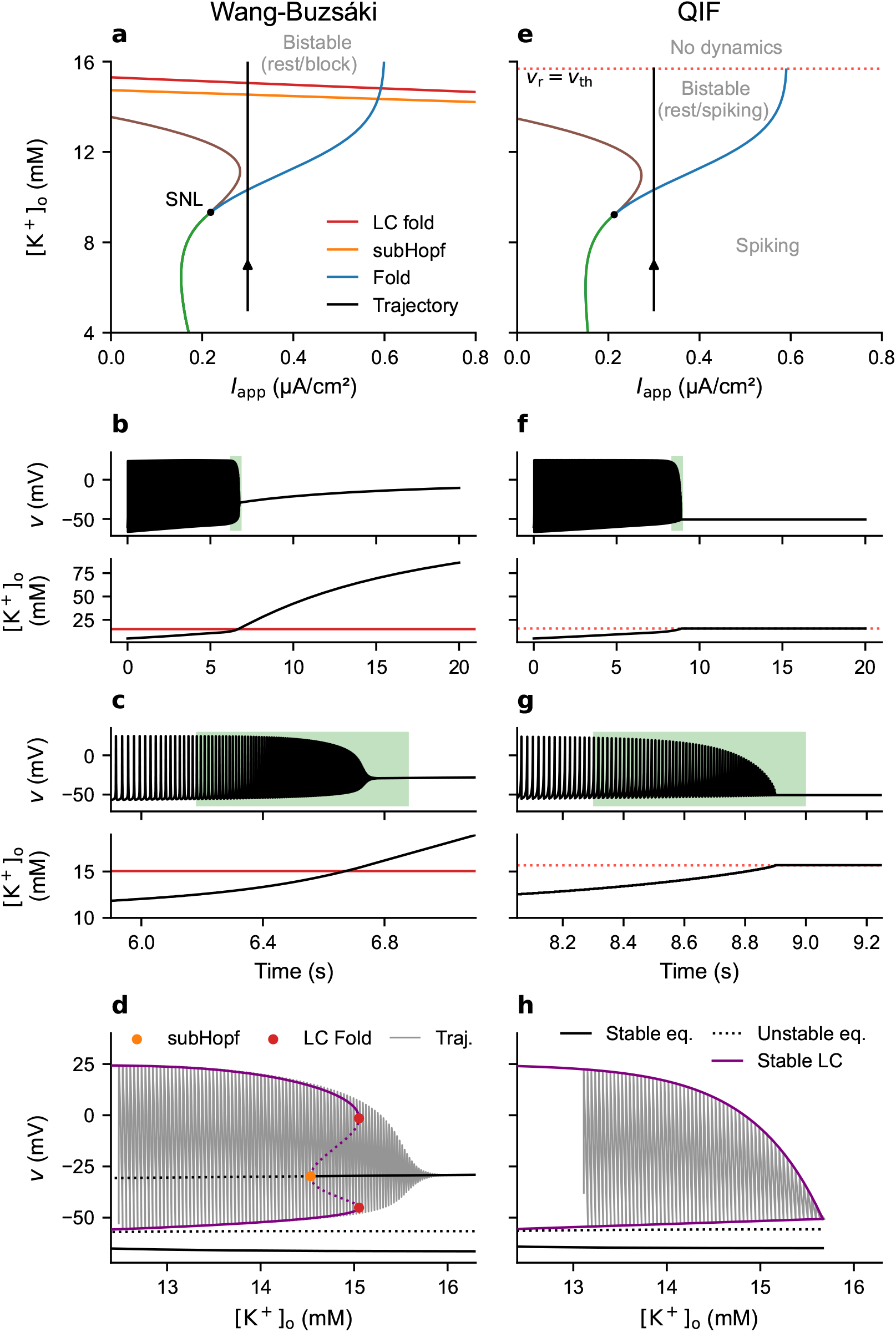
Depolarization block scenario (deterministic input: *I*_app_ = 0.3 µA cm^−2^). For the Wang-Buzsáki model: **(a)** Trajectory shown on the 2-parameter bifurcation diagram [K^+^]_o_ versus *I*_app_ of the fast subsystem. **(b)** Voltage and [K^+^]_o_ time traces. **(c)** Enlargement of the transition to depolarization block shown in panel (b). **(d)** Part of the trajectory highlighted in panels (b) and (c), superimposed onto the 1-parameter bifurcation diagram of the fast subsystem with respect to [K^+^]_o_. For the quadratic integrate-and-fire model: **(e-h)** Same as (a-d).

When stimulated with the same deterministic input, the integrate-and-fire version of the model produces a voltage trace very similar to the depolarization block observed in the Wang-Buzsáki model (Fig. 8f-g). However, unlike the two other scenarios, this behavior does not entirely rely on the same mechanism as in the original model. The path leading to the block is unchanged, but the transition to the block itself cannot occur via the same dynamical bifurcation, due to the design of the reduced model. Instead, we emulate it by placing the system in a refractory state as soon as, at larger [K^+^]_o_ values, the reset voltage meets the threshold value (Fig. 8e and h): the dynamics of both the voltage and [K^+^]_o_ are stopped, and their values remain fixed thereafter (see Fig. 8f and g). With this constraint, although the dynamics during the block is only sketchily captured, the desired outcome is for the neuron to cease spiking. The dynamics preceding the block, by contrast, is captured satisfactorily: the QIF voltage trace features both the reduction in spike amplitude and increase in firing frequency (Fig. 8g). These two effects are in agreement with the experimental recordings in [27]. As shown in Fig. S3f and h, the frequency of the fast subsystem limit cycles as a function of [K^+^]_o_ increases in two stages before the block. This pattern is a remnant of the situation with two homoclinic bifurcations at lower values of applied current (Fig. S3b and d).

### Reshaping of the phase-response curve

We now compare the phase-response curves (PRCs) of the two models. For a spiking neuron, the PRC describes how the phase of the cycle at which a perturbation is received affects the timing of the next action potential. It provides a link between the dynamics of single neurons and their collective behavior (e.g., synchronization) when connected in a network.

Fig. 9a shows the infinitesimal PRC (iPRC, i.e., for infinitesimal input amplitude) of the Wang-Buzsáki model near spike onset for different potassium concentrations. Like when the SNL is reached via other biophysical parameters (e.g., see [30] for the capacitance), we can see a transition from almost symmetric at low [K^+^]_o_ to left-tilted PRCs after the SNL bifurcation. Near spike onset, this alteration of the iPRC occurs suddenly at the SNL: the curves for [K^+^]_o_ = [K^+^]_o,SNL_ − 4 mM and [K^+^]_o_ = [K^+^]_o,SNL_ − 2 mM are still nearly identical.

**FIG. 9:**
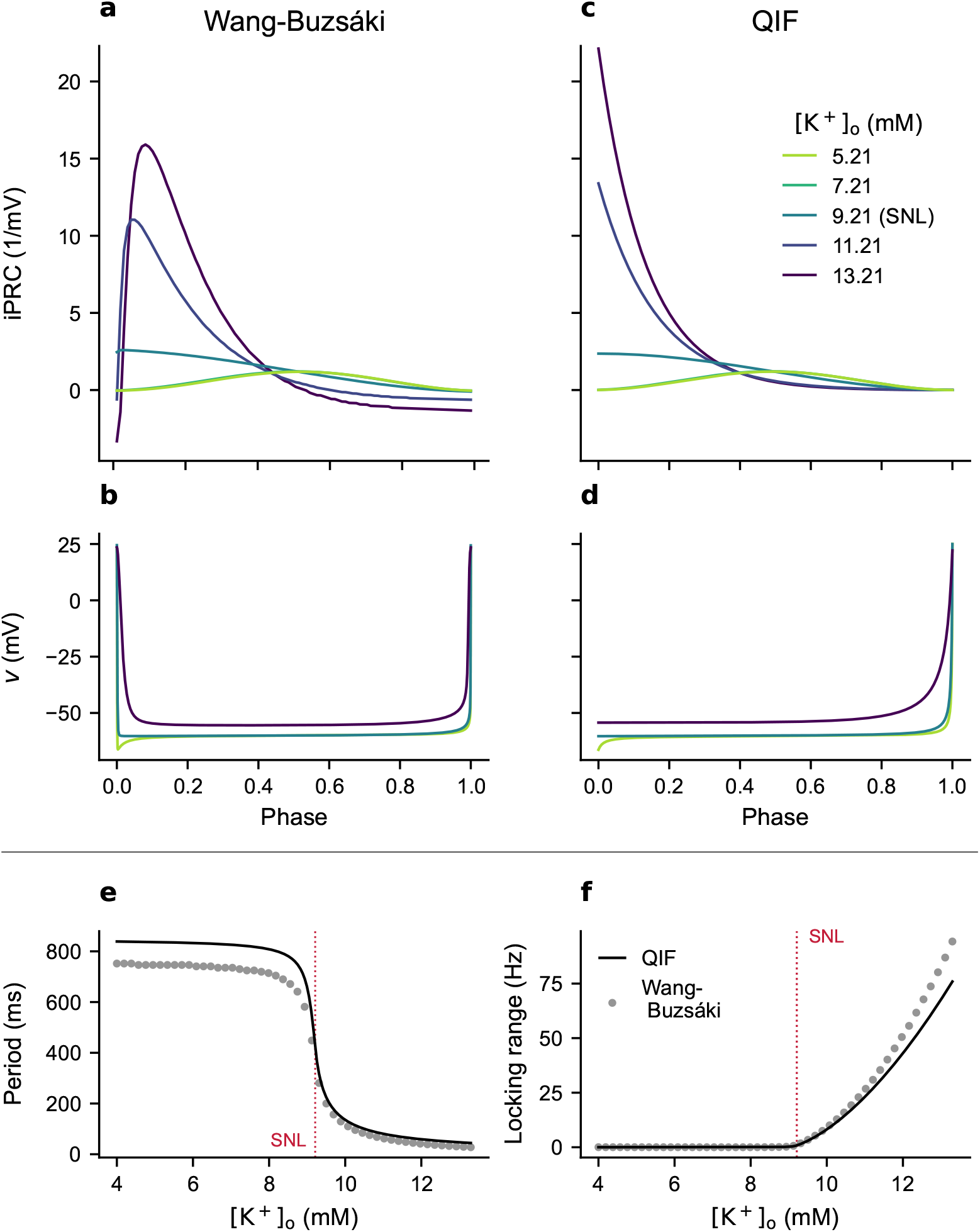
Effect of [K^+^]_o_ on the phase response curve and ability to synchronize near spike onset 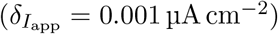. **(a)** iPRCs for the Wang-Buzsáki model (direct method). Note that the curve for [K^+^]_o_ = 7.21 mM lies beneath the curve for [K^+^]_o_ = 5.21 mM; the same holds for panel (c). **(b)** Spike voltage traces for the Wang-Buzsáki model. **(c)** iPRCs for the integrate-and-fire model (analytically, see “Methods”). **(d)** Spike voltage traces for the integrate-and-fire model. **(e)** Period of the limit cycles. **(f)** Locking range, assuming pulse coupling with a voltage perturbation of 0.15 µV. For the computation of the coupling function, pulses were normalized following Weerdmeester et al. [31].

The PRC reshaping at the SNL is well captured by the QIF model (Fig. 9c). In fact, this simplified model offers an intuitive understanding of how an increase in [K^+^]_o_ deforms the iPRC. Since we are dealing here with a one-dimensional system, the iPRC is simply the inverse of the derivative of the spike voltage trace as a function of the phase. Flatter slopes therefore correspond to larger iPRC values. At low [K^+^]_o_, the inflection point of the spike is near phase (Fig. S5b), resulting in an almost symmetric iPRC (Fig. S5a). The effects of larger [K^+^]_o_ are mainly expressed in the QIF model by a larger reset voltage, which results in skipping the beginning of the spike cycle (see Fig. 9d or compare panels b and d in Fig. S5). Increasing the reset is thus equivalent to retaining only a scaled version of the iPRC segment corresponding to late phases (see Fig. 9c or compare panels a and c in Fig. S5), resulting in more left-tilted iPRCs.

As pointed out in [30], the SNL bifurcation coincides with a reduction of the period at a given distance 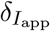 from spike onset (i.e., sharper f-I curves). This period reduction is captured by the QIF version of the model (Fig. 9e).

Note that, at large [K^+^]_o_, iPRCs contain a negative region (corresponding to phase delay) early in phase. This feature is not captured in the QIF model, related to the fact that QIF models do not explicitly describe membrane repolarization. Their iPRCs therefore cannot contain negative regions.

The changes associated with the SNL bifurcation directly affect the collective behavior of neurons in a network. Based on the theory of weakly coupled oscillators [32], and given a specific network configuration, the iPRC can be used to predict the neurons’ ability to synchronize. For example, in the case of two weakly pulse-coupled neurons (where each spike induces a small, instantaneous voltage jump in the postsynaptic neuron), the coupling function is a scaled version of the iPRC, and its odd part governs how the phase difference between the two neurons evolves over time [9, 31, 33]. When the uncoupled firing frequency differs between the two neurons, the locking range, i.e., the maximal frequency detuning such that there exists a stable fixed point for the dynamics of the phase difference, is twice the amplitude of the odd part of the coupling function [9]. Fig. 9f shows the locking range as a function of extracellular potassium concentration for a voltage perturbation amplitude of 0.15 µV. For both models, the locking range abruptly increases at the SNL, indicating enhanced synchronization ability. In more complex networks, the analysis must be adapted to the particular network architecture (e.g., network topology and synaptic coupling), but the PRC can still serve to predict aspects of the network state [34].

### Extracellular potassium influences neuronal synchronization in a network

We now investigate the consequences of the changes introduced by the SNL bifurcation at the network scale. In particular, for tonically spiking neurons, we expect that the left-tilted PRCs at larger extracellular potassium concentrations induce in-phase synchronization of inhibitory networks [11]. We test this prediction in a basic network configuration. For this, we consider a network of 100 all-to-all pulse-coupled inhibitory neurons. We then impose on all these neurons a common extracellular potassium trace, simulating, for instance, an external wave reaching the network (Fig. 10a). With both models, the neurons of the network, which were previously out of synchronization (Fig. 10c and f), become in-phase synchronized upon arrival of this wave (Fig. 10d and g).

**FIG. 10:**
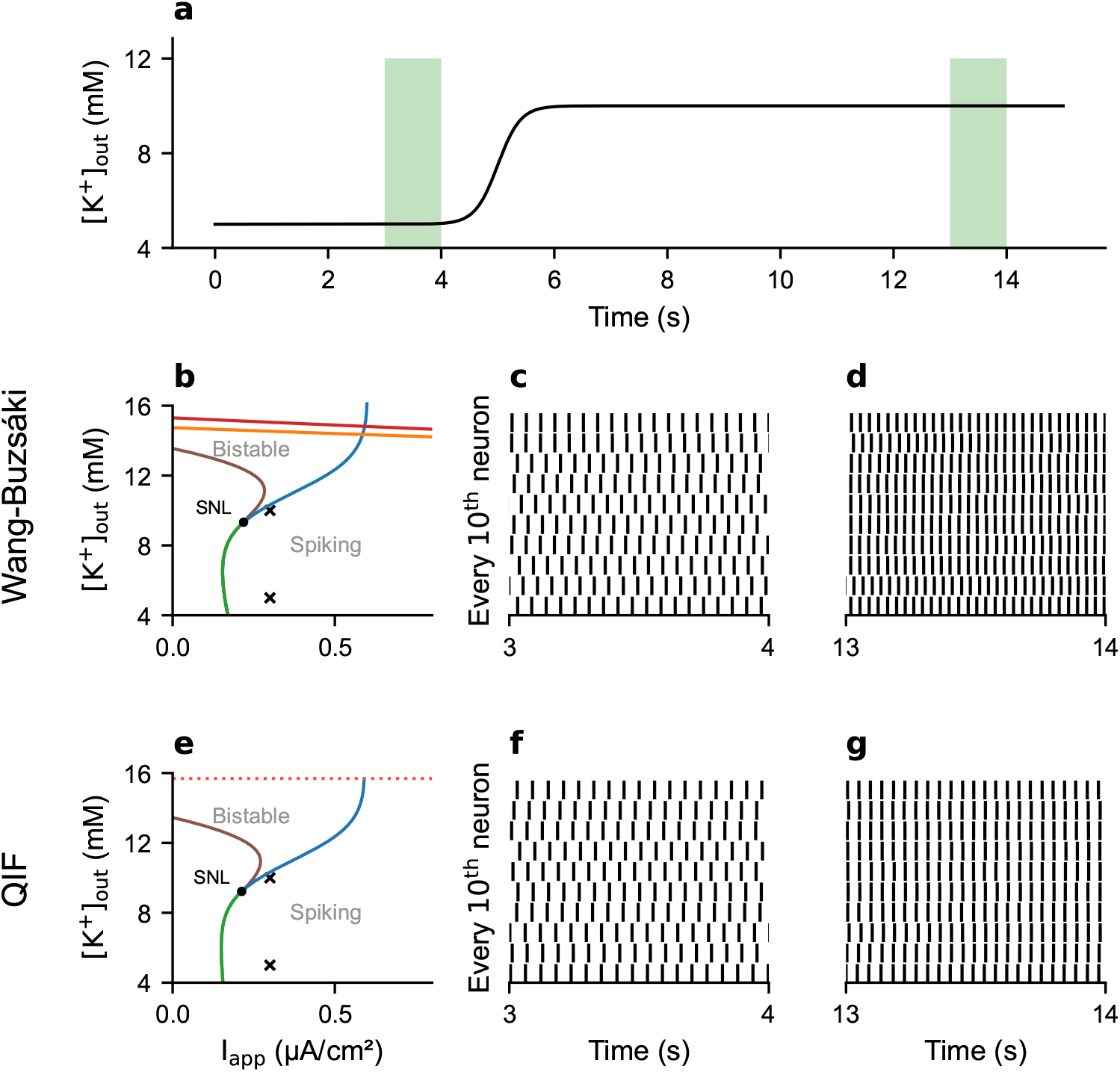
Response of a network of 100 neurons to an external potassium wave. We considered an all-to-all connectivity scenario, with weak inhibitory pulse-coupling (*δ*_*v*_ = −0.000 15 mV). We used a noisy input (see “Methods”), common to all neurons, with mean *I*_app_ = 0.25 µA cm^−2^. **(a)** Incoming potassium wave, common to all neurons. **(b)** 2-parameter bifurcation diagram [K^+^]_o_ versus *I*_app_ of the fast subsystem for the Wang-Buzsáki model. The crosses indicate the initial and final potassium concentrations. **(c-d)** Raster plot for 10 among the 100 neurons, during the third and 13^th^ seconds, respectively. **(e-g)** Same as (b-d), for the QIF version of the model.

Disentangling the respective contributions of PRC asymmetry and frequency fluctuations to the effect of the SNL transition on synchronization requires further investigation. Furthermore, we deliberately chose [K^+^]_o_ values such that the neurons remain in the tonic spiking region (Fig. 10b and e). Indeed, in the bistable region, where only a fraction of the neurons may be active, the PRC is not a reliable predictor of network behavior. A detailed study of the network dynamics is beyond the scope of this article (see, for example, [35]).

## Discussion

Neuronal dynamics can influence network behavior, yet the inclusion of biophysical detail in larger network models both challenges computational efficiency and reduces model generality due to the number of additional parameters introduced. Commonly, neuron models are fitted to datasets from a single experimental setting, and one best-fit parameter set is determined. These models may fail to generalize when parameters not varied in the experiment are altered in the model. Here, we derive models that capture the impact of a biophysical parameter across a physiologically relevant range by fitting bifurcation diagrams rather than voltage traces tied to specific recording conditions. The resulting models capture the dynamical transitions across the fitted parameter range, at the cost of reduced precision at individual points in parameter space.

At the scale of a single neuron, we have shown that the phenomenological model accurately reproduces various activity patterns from the original model, without needing to adjust parameter values to each scenario. At the network level, preliminary simulations highlight the important role that extracellular potassium concentration plays in synchronization.

Specifically, we have here shown how to reduce a class I conductance-based model with dynamic potassium concentration to a quadratic integrate-and-fire model. Depending on the concentration of extracellular potassium and its dynamics, single-cell excitability in this model can vary between the SNIC and HOM types, capturing the regime transitions observed in the conductance-based neuron model. The effect of potassium on the bifurcation structure in the original, conductance-based model is in the reduced QIF model version implemented via a change in the reset voltage. Note that this approach is not limited to potassium; as illustrated for temperature in Fig. S2, it can be extended to other parameters known to induce an SNL transition.

### A phenomenological neuron model capturing potassium dynamics

Phenomenological models that capture biophysical aspects are particularly valuable for exploring aspects at the network scale through systematic simulations. For example, phenomenological models of spike-frequency adaptation have been used to study network synchronization [2, 36]. In the same vein, our work formulates a simple model of potassium concentration dependence. It provides a minimal description of a dynamics that would typically require a more detailed, higher-dimensional system.

Extracellular potassium concentration is a particularly pertinent variable to retain in simplified neuron models. Typically, models assumed constant ion concentrations. This is motivated by the fact that the amount of relocated ions per action potential may have a negligible effect on the overall concentrations [37, 38], and that concentrations are regulated by mechanisms involving the Na^+^/K^+^-ATPase, cotransporters, and glial cells [39, 40]. Nonetheless, in the last decades numerous studies have focused on modeling their dynamics [13, 14, 16, 21, 27, 29, 41–47]. Indeed, experimental studies suggest activity-dependent fluctuations in concentration, in particular transient accumulations of extracellular potassium reaching several millimolars, in case of intense activity or following sensory stimulation [12, 39, 40, 45, 48, 49]. Such changes are expected to be particularly significant locally or in restricted spaces [39, 40]. Moreover, drastic increases in extracellular potassium are known to occur during pathophysiological events such as spreading depolarizations and epileptic seizures, as well as during hypoxia [39]. Both experimental evidence [21, 49–51] and results based on biophysical neuron models (e.g., [13, 14, 16, 21, 27, 29, 41, 47]) indicate that the weakening of the potassium driving force, resulting from activity-mediated extracellular build-up, in turn qualitatively alters neuronal excitability, at both moderate and large [K^+^]_o_ elevations. This motivates the need for simple models capturing potassium dynamics to investigate the consequences of changes in neurons’ intrinsic excitability at the network scale.

There exist other minimalist models with potassium dynamics, mainly in the context of epilepsy [52–54]. The single neuron model by Depannemaecker et al. [52] reproduces a wide range of firing patterns, including seizure-like events, sustained ictal activity, and depolarization block, while being relatively simple. It remains, nevertheless, at a higher level of complexity compared to the present model, with its two-dimensional fast subsystem of type Hodgkin-Huxley and its two slow variables. The “epileptor-2” population model [53] by Chizhov et al. is also four-dimensional, with the advantage of explicitly modeling sodium dynamics. It can successfully simulate interictal and ictal discharges in high potassium conditions. However, the choice of coupling between potassium and voltage dynamics in this model does not permit onset bifurcation switches in individual neurons. In a recent study, Depannemaecker et al. propose an extension of the adaptive exponential integrate-and-fire model (AdEx) that accounts for impaired ionic gradients [54], by introducing a third, slow, variable.

### Modeling the critical transitions in parameter space

The question of “what is a good model” has been debated extensively in the epistemological literature and many arguments in favor of simple leaky integrate-and-fire models have been presented [55]. For example, if well fitted, they have high predictive power for the voltage state variable. However, if one includes model predictions not only in observable state variables, but also for changed biophysical parameter sets, preserving the relevant bifurcation structure can improve robustness.

The idea that neuronal models should be understood through their bifurcation structure is not new in computational neuroscience. Beginning with FitzHugh’s phase-plane analysis and reduced models [56, 57], the emphasis gradually shifted from reproducing every ionic current to capturing the essential dynamical mechanisms underlying neuronal behavior. Building on this perspective, Rinzel and Ermentrout demonstrated that classes of neural excitability correspond directly to bifurcation types [58]. Rinzel also introduced slow–fast analysis to explain bursting dynamics [59]. Bertram et al. extended this viewpoint by classifying bursting according to the unfolding of codimension-2 or higher bifurcations of the fast subsystem [60], a framework that was subsequently broadened by Golubitsky et al. [61]. Izhikevich further organized this landscape by classifying bursters according to the pair of codimension-1 bifurcations that bound their silent and active phases [8, 62]. This framework continues to be developed and extended [30, 63, 64], making the construction of reduced models that match the relevant bifurcation diagram a natural next step.

Our work, therefore, presents a strategy to derive models that capture the dependency of the system on physiological quantities by matching the relevant bifurcation diagram. For a related approach in a broader context, see [65]. This way of constructing phenomenological models is a real alternative to approaches based on fitting spike trains, in particular because neurons are not stationary devices, but subject to constant change in their environmental parameters. Sources of variability include temperature [9, 66, 67], ionic concentrations [16, 21, 68] and neuromodulators [69]. These factors may directly depend on the recent history of activation [21, 70]. In some cases, their changes constitute a homeostatic response to other changes, helping to maintain the neurons’ properties in the face of a varying environment [71]. We argue that, when known, fitting the bifurcation diagram with respect to chosen parameters is a judicious way to obtain models tailored to the major source of instability in a given context, as we have done here with the extracellular potassium concentration. Fitting specific spike trains may result in models whose validity is largely restricted to the conditions of their construction. In contrast, fitting a bifurcation diagram may not only provide better robustness to changing conditions but also allow for the study of how these changes themselves slowly drive neuronal activity between different regimes.

In this study, we use as a target the bifurcation diagram of a conductance-based neuron model. This target is only indirectly constrained by experimental data, through gating functions fitted to voltage-clamp recordings. Ideally, the target bifurcation diagram would instead be obtained directly from experimental measurements. Estimating bifurcation diagrams from experiments, however, remains challenging. Some techniques exist to approximate branches of stable and unstable steady states, and branches of stable limit cycles can be approximated using slowly ramped current-clamp protocols [72]. In practice, however, many factors vary during recordings, and generating reliable diagrams would require tightly controlled experimental repetitions in various conditions, particularly for reconstructing two-dimensional bifurcation diagrams. Developing such protocols is beyond the scope of the present study but represents an important direction for future work.

There exist canonical phenomenological models that, thanks to their bifurcation landscape, can recapitulate the main features of neural excitability (see, for example, [73], or [63], which is based on the unfolding of a higher-dimensional bifurcation). However, they are formulated in terms of abstract variables. In some situations, having a model that contains specific bifurcations is not sufficient; their arrangement in terms of biophysical dimensions also matters. For example, the fold/hom bursting mechanism that we studied in the first scenario cannot occur when the unfolding of the SNL bifurcation is aligned with the [K^+^]_o_ axis (see [13] for more details). Starting from a canonical model and adapting it to match desired bifurcation branches, as we did here with the QIF model, has the advantage of preserving important quantitative components.

### Possible model extensions

#### Sodium dynamics

In this work, we restricted ourselves to potassium dynamics, although sodium concentrations are also directly affected by neuronal activity. This choice was made because potassium is the main player in activity-induced SNL transitions and switches in excitability class, while sodium is rather involved in spike amplitude reduction [21]. The engagement of the Na^+^/K^+^-ATPase to restore ionic gradients is accounted for in our model by the potassium dependence of the pump current. In [13], it is shown that sodium dynamics is not crucial for the bursting mechanism we studied in the first scenario.

Taking into account the dynamics of the two ion species, for example by tracking extracellular potassium and intracellular sodium concentrations, remains the most precise option [53, 74]. To avoid adding an extra variable, one could rely on electroneutrality to express one concentration as a function of the other [27, 28, 52]. Applying this strategy to our work is an interesting direction for refinement.

#### Potassium diffusion and spatial network models

A natural extension of the present model is its use in spatially structured networks in which extracellular potassium can diffuse between local compartments. The reduced model is particularly suited to this purpose because it retains the potassium-dependent transitions in neuronal excitability while remaining computationally inexpensive enough for large network simulations. At the single-neuron level, Fig. S4 already illustrates that adding a simple potassium diffusion term can stabilize extracellular potassium and thereby qualitatively alter the resulting firing regime. In a spatial network, allowing potassium released by active neurons to spread to neighboring extracellular compartments would introduce an additional, non-synaptic interaction between neurons, through which local activity can modify the excitability of nearby cells. Such models could be used to investigate how synaptic and potassium-mediated diffusive coupling interact to shape synchronization, spatial propagation of activity, and transitions between excitability regimes (see [75, 76]).

#### Transition to depolarization block

Finally, the way in which depolarization block is obtained in the reduced model (Scenario 3) is not ideal. Instead of being mediated by a supercritical Hopf bifurcation, or a subcritical Hopf coupled to a fold of limit cycles, as in conductance-based models, it was artificially achieved by silencing the dynamics when the reset and threshold voltages meet. Indeed, the fast subsystem of the reduced model, of type quadratic integrate-and-fire, cannot exhibit Hopf bifurcations. Instead, one would need to start from a two-dimensional fast subsystem, like the AdEx model. This would not only allow a more accurate modeling of the transition to the block but also open the possibility of studying the recovery from this pathological state.

## Conclusion

In conclusion, we affirm the importance of phenomenological models that capture key underlying biophysical mechanisms. We contribute a model of potassium concentration dependence, readily suitable for integration into large networks. Moreover, our approach, based on the fitting of a bifurcation structure, provides a generalizable framework for deriving similar models for other relevant modulatory variables.

## Methods

### Extended Wang-Buzsáki model [13]

#### Model definition

The model obeys the system of differential equations

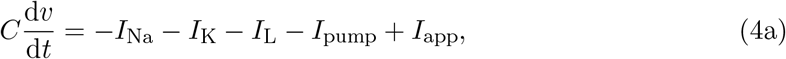

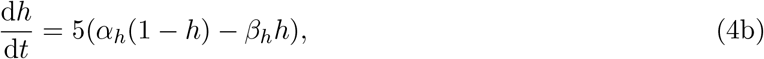

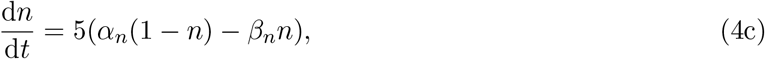

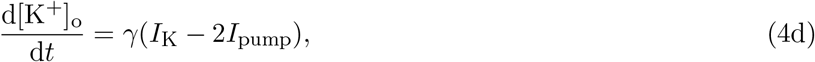

with currents

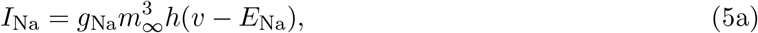

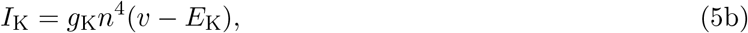

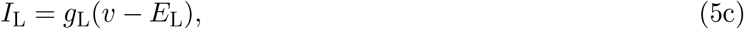

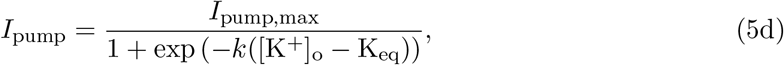

and gating functions

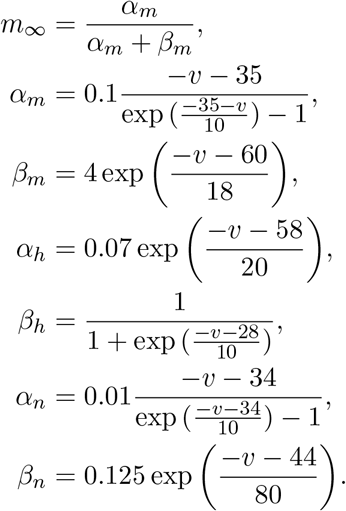

The reversal potential for the potassium evolves with [K^+^]_o_ according to

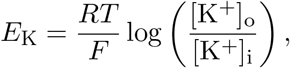

where *R* = 8314.4 mJ K^−1^ mol^−1^ is the ideal gas constant, *F* = 96 484.6 C mol^−1^ the Faraday’s constant and *T* = 310 K the temperature. The variables and parameter values are listed in Tables III and IV.

**TABLE III:**
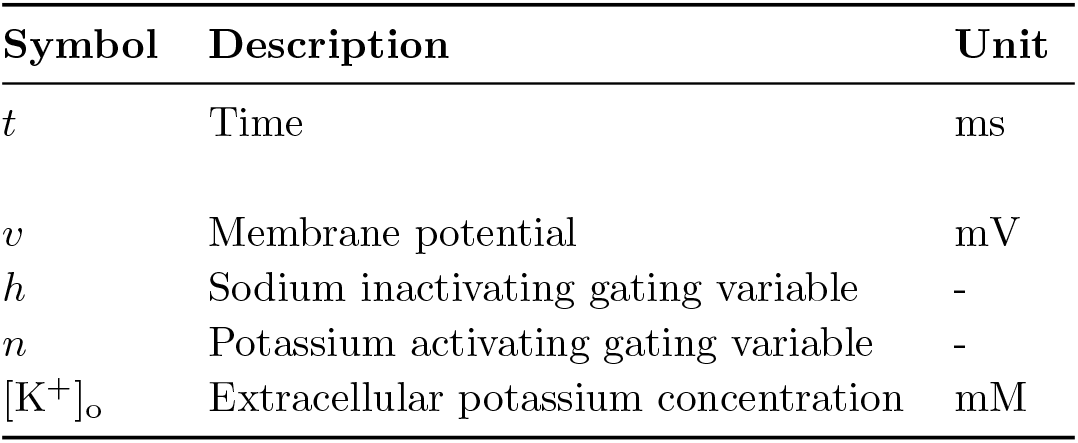
Variables of the extended Wang-Buzsáki model [13] given in Eq. (4).

| Symbol | Description | Unit |
| --- | --- | --- |
| $t$ | Time | ms |
| $v$ | Membrane potential | mV |
| $h$ | Sodium inactivating gating variable | - |
| $n$ | Potassium activating gating variable | - |
| $[K^+]_o$ | Extracellular potassium concentration | mM |

**TABLE IV:** Parameter values for the extended Wang-Buzsáki model [13] given in Eq. (4).

| Symbol | Description | Value | Unit | Source |
| --- | --- | --- | --- | --- |
| $I_{app}$ | Applied current | Varied | $\mu A cm^{-2}$ | |
| $C$ | Membrane capacitance density | 1 | $\mu F cm^{-2}$ | [13, 15] |
| $g_L$ | Leak conductance | 0.1 | $mS cm^{-2}$ | [13, 15] |
| $E_L$ | Leak reversal potential | -65 | mV | [13, 15] |
| $g_{Na}$ | $Na^+$ maximal conductance | 35 | $mS cm^{-2}$ | [13, 15] |
| $E_{Na}$ | $Na^+$ reversal potential | 55 | mV | [13, 15] |
| $g_K$ | $K^+$ maximal conductance | 9 | $mS cm^{-2}$ | [13, 15] |
| $[K^+]_i$ | Intracellular potassium concentration | 140 | mM | [13, 77] |
| $I_{pump,max}$ | Maximal pump current | 0.5 | $\mu A cm^{-2}$ | |
| $k$ | Pump activation steepness | 0.9 | $mM^{-1}$ | [13] |
| $K_{eq}$ | $[K^+]_o$ for half pump activation | 11 | mM | [13] |
| $\gamma$ | Conversion factor | $4.1457 \times 10^{-4}$ | $mM ms^{-1}/(\mu A cm^{-2})$ | [13, 77] |

#### Slow dynamics

The standard slow subsystem of System (4) is given by

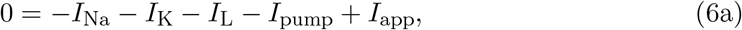

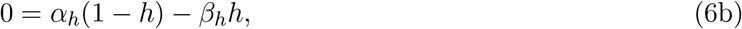

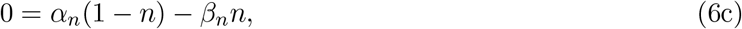

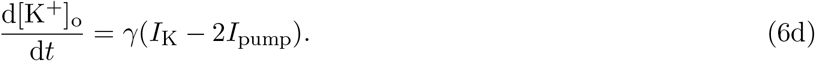

As in [13], for [K^+^]_o_ values such that there is a stable limit cycle in the fast subsystem, we define the averaged slow subsystem of System (4) as

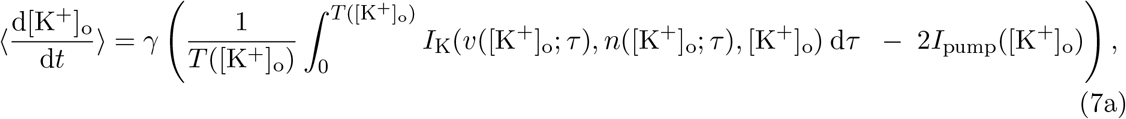

where *T* is the period of the limit cycle.

### Fast subsystem of the QIF model (Eqs. (3a) and (3c))

#### Frequency of the limit cycles

Let 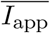= *c*(*I*_app_ − *I*_SN,0_ − *I*_pump_). If 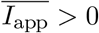 (tonic spiking region), the period of the limit cycles is given by:

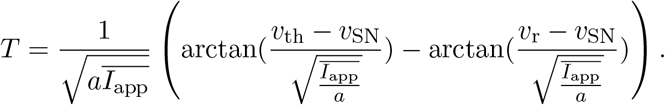

If *v*_r_ *> v*_SN_ and 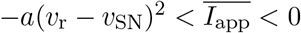 (bistable region):

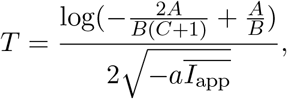

with

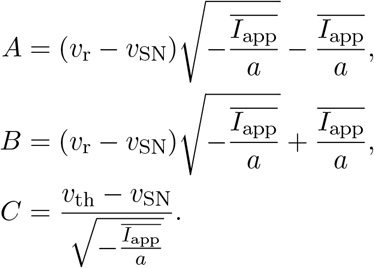

Infinitesimal phase-response curve (iPRC)

Let. 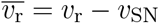 If 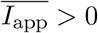:

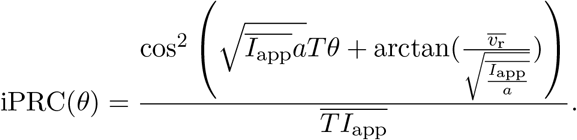

If *v*_r_ *> v*_SN_ and 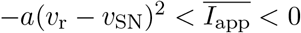 :

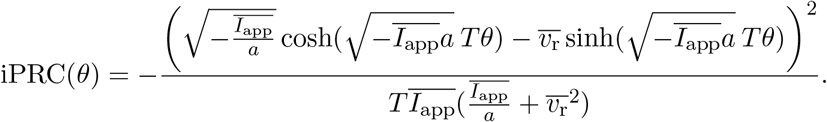

### Noisy input

To introduce noise in the applied current, we replace it with:

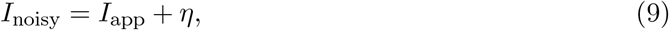

where *η*_*t*_ is an Ornstein-Uhlenbeck (OU) process defined by

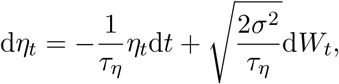

*W*_*t*_ being the Wiener process. The OU process can be interpreted as a smoothly low-pass filtered white noise. In our simulations, we used a correlation time *τ*_*η*_ = 20 ms and a standard deviation *σ* = 0.05 µA cm^−2^.

### Computational resources

To derive the parameters *I*_SN,0_, *v*_SN_, *a* and *v*_r_ of the QIF model (1), we performed symbolic computations with the Python library Sympy [78]. In the case of *v*_SN_, we additionally used the numerical solver nsolve.

For numerical continuation, we used the software AUTO-07P [79].

Simulations of the neuron models were carried out with the Python package brian2 [80].

## Acknowledgments

This project has received funding from the European Research Council (ERC) under the Union’s Horizon 2020 research and innovation program (grant agreement no 864243), from the Einstein Foundation Berlin (to SS, EP-2021-621), and from the German Research Council (CRC/TRR 384). The funders had no role in study design, data collection and analysis, decision to publish, or preparation of the manuscript.

## Supplementary figures

**FIG. S1:**
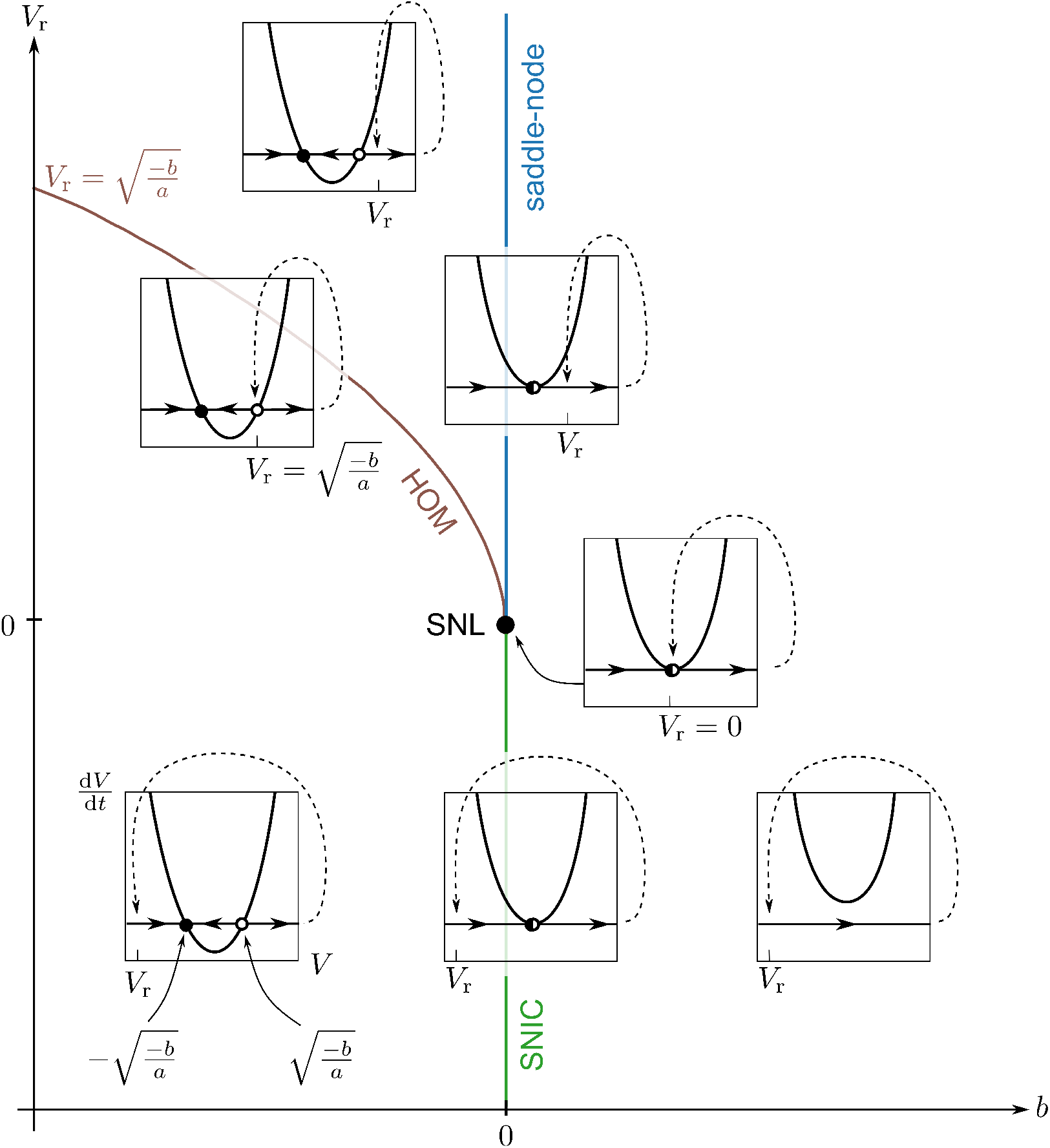
Bifurcation diagram of the classical QIF model, defined by: 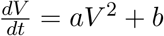 (with *a >* 0); if *V > V*_th_, then *V* ← *V*_r_. When *b* is negative, the system has a stable fixed point at 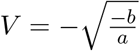 and an unstable fixed point at 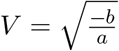. They disappear via a saddle-node bifurcation at *b* = 0. When the reset voltage *V*_r_ is negative, periodic solutions emerge via a SNIC bifurcation at *b* = 0. When *V*_r_ is positive, periodic solutions are instead created via a homoclinic bifurcation, at 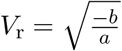. In this case, for 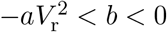, the system exhibits bistability between the stable limit cycle and the stable fixed point. The transition from SNIC spike onset to homoclinic spike onset is mediated by a SNL bifurcation at *V*_r_ = *b* = 0 (i.e., when the voltage is reset to the saddle-node point), a codimension-2 bifurcation which acts here as an organizing center. Adapted from Figure 8.3 in [8].

**FIG. S2:**
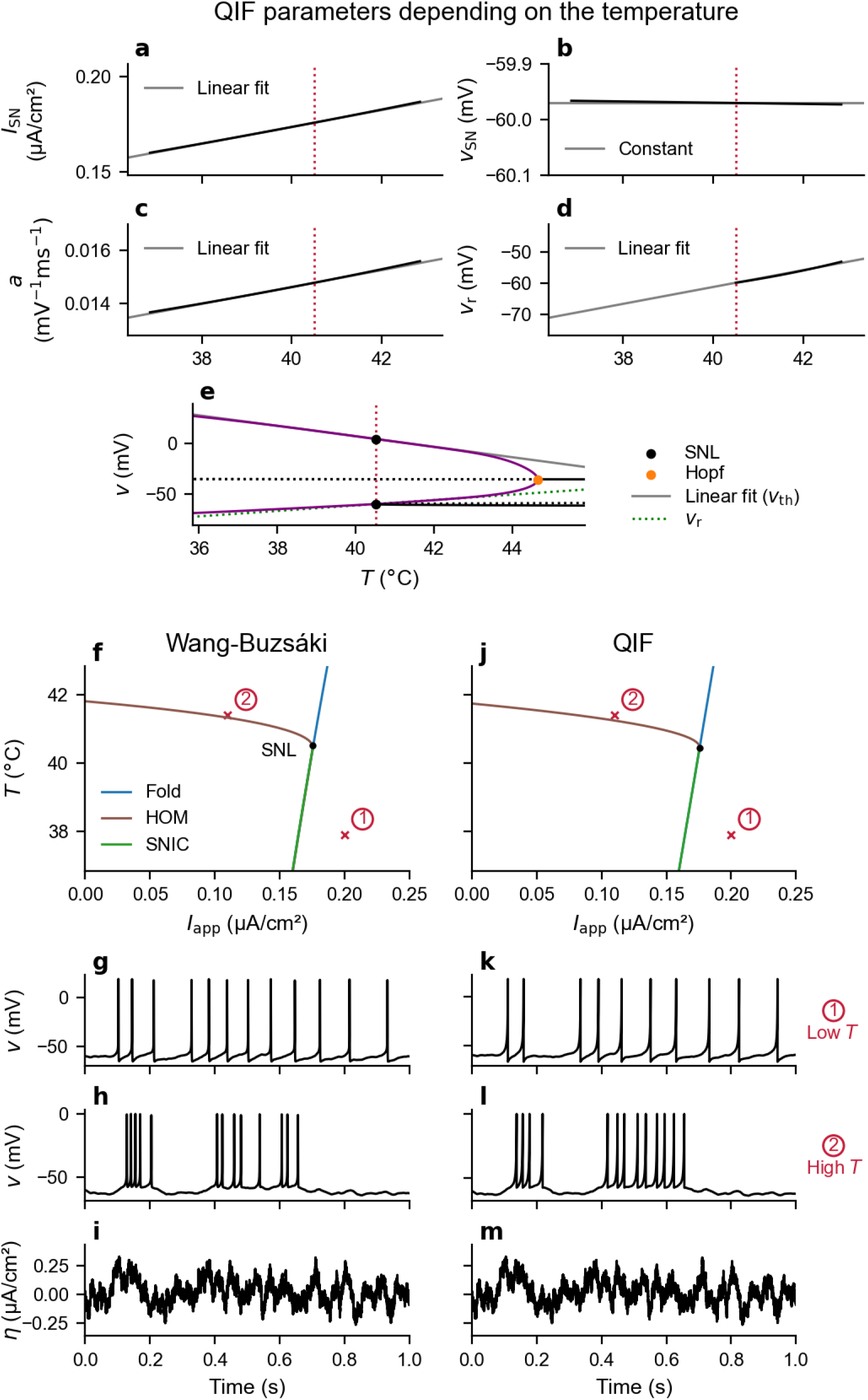
Illustration of the reduction procedure, here applied to derive a QIF model that captures dependence on temperature instead of potassium concentration, motivated by Hesse et al. [9]. The initial conductance-based model is the Wang–Buzsáki model as in [9]: [K^+^]_o_ is constant, there is no pump current, and temperature dependence is introduced via the Nernst equation and Q_10_ coefficients. The resulting QIF model reproduces the transition induced by temperature rise reported in Figure 3 of [9], characterized by intermittently interrupted firing in response to noisy input. **(a-e)** QIF parameters derivation: *I*_SN_ (a), *v*_SN_ (b), *a* (c), *v*_r_ (d) and *v*_th_ (e).**(f-j)** Two-parameter bifurcation diagram temperature *T* versus applied current *I*_app_, for the initial (f) and QIF (j) models. **(g-i)** Voltage response to a noisy input current *I*_noisy_ (Eq. (9) in “Methods”, with *τ*_*η*_ = 0.01 s and *σ* = 0.1 µA cm^−2^) at low (g) or high (h) temperature, in the initial model. The mean applied current *I*_app_ is chosen smaller in the HOM case (high temperature) than in the SNIC case, to obtain a similar number of spikes per second (see red crosses in panels (f) and (j)). The time course of *η* is shown in panel (i). **(k-m)** Same as (g-i), in the QIF model.

**FIG. S3:**
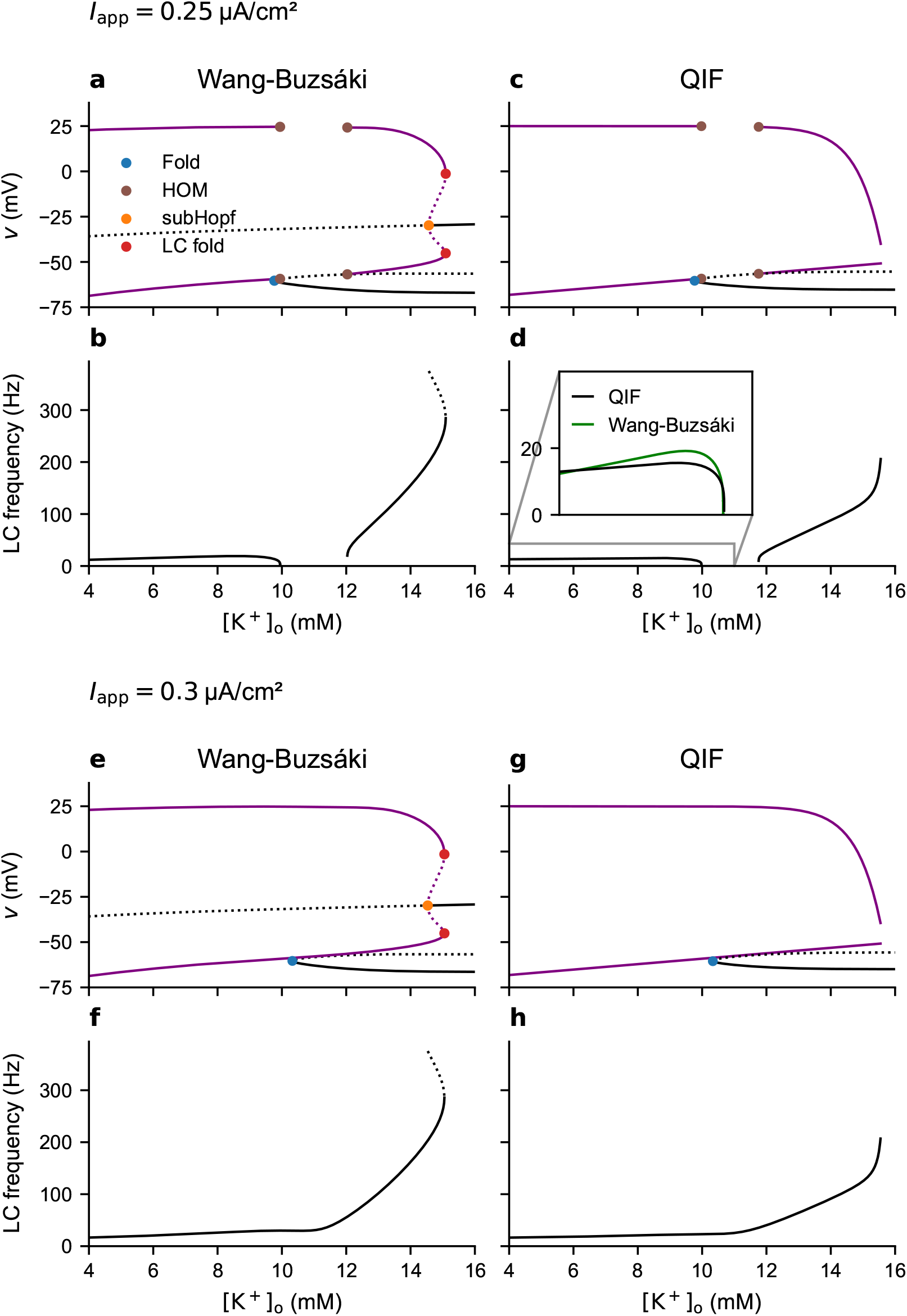
Bifurcation diagrams with respect to [K^+^]_o_ and frequency of the limit cycles when *I*_app_ = 0.25 µA cm^−2^, for the Wang-Buzsáki **(a**,**b)** and integrate-and-fire **(c**,**d)** models. **(e-h)** Same as **(a-d)**, when *I*_app_ = 0.3 µA cm^−2^.

**FIG. S4:**
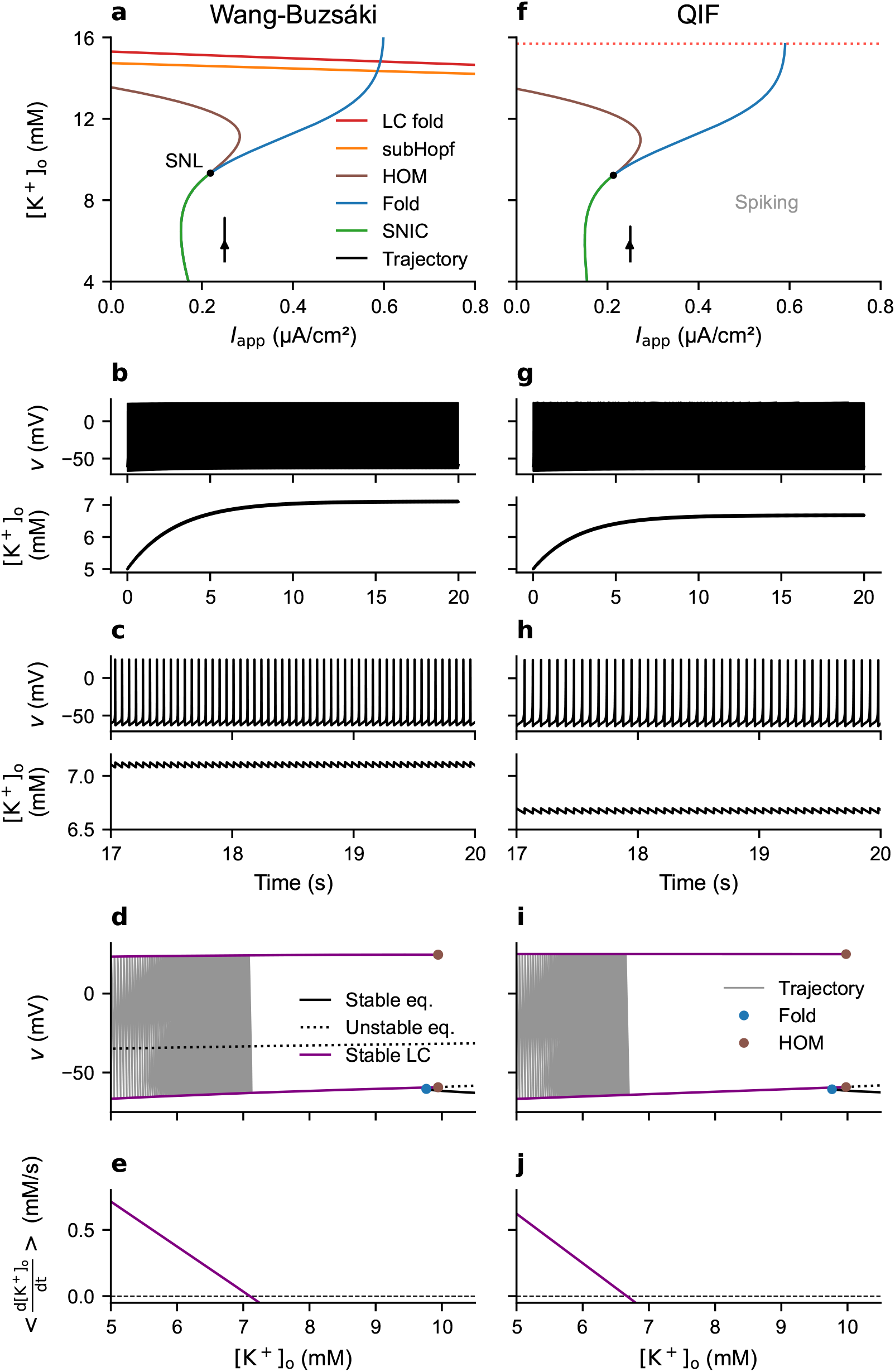
Tonic firing scenario, when adding a diffusion term *I*_diff_ = *D*([K^+^]_o_ − *K*_bath_) to the potassium dynamics, with diffusion coefficient *D* = 0.4 Hz and bath concentration *K*_bath_ = 5 mM. The initial conditions and other parameter values are unchanged compared to the fold/homoclinic bursting scenario. For the Wang-Buzsáki model: **(a)** Trajectory shown on the 2-parameter bifurcation diagram [K^+^]_o_ versus *I*_app_ of the fast subsystem. **(b)** Voltage and [K^+^]_o_ time traces. **(c)** Enlargement of (b). **(d)** Trajectory superimposed onto the 1-parameter bifurcation diagram with respect to [K^+^]_o_. **(e)** Time derivative of the averaged slow subsystem. **(f-j)** Same as (a-e), for the integrate-and-fire version.

**FIG. S5:**
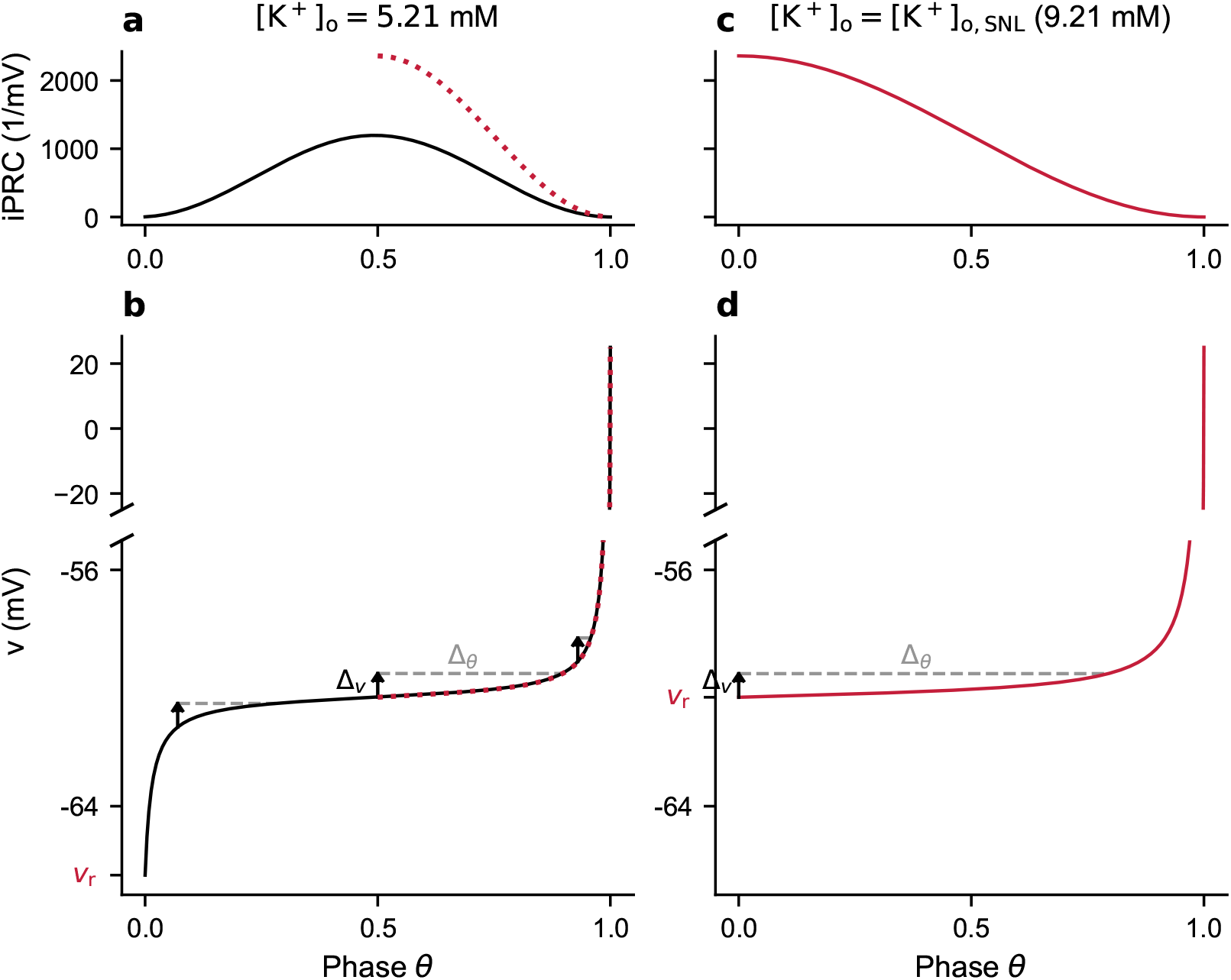
Schematic representation of the relation between the spike voltage trace and iPRC in the QIF model, a one-dimensional model. (**a-b**) iPRC near spike onset 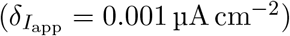 and spike voltage trace when [K^+^]_o_ = 5.21 mM (physiological). The iPRC is largest when the derivative of the solution is smallest. (**c-d**) iPRC near spike onset (same 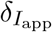) and spike voltage trace at the SNL ([K^+^]_o_ = 9.21 mM), i.e., when the reset is at the inflection point.

## Notes

### Competing Interest Statement

The authors have declared no competing interest.

### Summary of Updates

General algorithm added (Table I); second example with temperature instead of potassium concentration added (Fig. S2); Discussion expanded to better position our work within the literature on the role of bifurcation structure in neuronal dynamics; figure added for readers unfamiliar with the QIF model (Fig. S1); scope regarding comparison with experimental data clarified (Intro., Discussion).

